# What can 1.8 billion regressions tell us about the pressures shaping high-level visual representation in brains and machines?

**DOI:** 10.1101/2022.03.28.485868

**Authors:** Colin Conwell, Jacob S. Prince, Kendrick N. Kay, George A. Alvarez, Talia Konkle

## Abstract

The rapid development and open-source release of highly performant computer vision models offers new potential for examining how different inductive biases impact representation learning and emergent alignment with the high-level human ventral visual system. Here, we assess a diverse set of 224 models, curated to enable controlled comparison of different model properties, testing their brain predictivity using large-scale functional magnetic resonance imaging data. We find that models with qualitatively different architectures (e.g. CNNs versus Transformers) and markedly different task objectives (e.g. purely visual contrastive learning versus vision-language alignment) achieve near equivalent degrees of brain predictivity, when other factors are held constant. Instead, variation across model visual training diets yields the largest, most consistent effect on emergent brain predictivity. Overarching model properties commonly suspected to increase brain predictivity (e.g. greater effective dimensionality; learnable parameter count) were not robust indicators across this more extensive survey. We highlight that standard model-to-brain linear re-weighting methods may be too flexible, as most performant models have very similar brain-predictivity scores, despite significant variation in their underlying representations. Broadly, our findings point to the importance of visual diet, challenge common assumptions about the methods used to link models to brains, and more concretely outline future directions for leveraging the full diversity of existing open-source models as tools to probe the common computational principles underlying biological and artificial visual systems.

## Introduction

The biological visual system transforms patterned light along a hierarchical series of processing stages into a useful visual format, capable of supporting object recognition (DiCarlo et al., 2012). Visual neuroscientists have made significant progress in understanding the nature of the tuning in early areas like V1 (Hubel and Wiesel, 1968), as well as in crafting normative computational accounts of the ecological and biological conditions under which such tuning might emerge (e.g. sparse coding over natural image statistics; Olshausen et al., 1995). However, that same computational clarity has been lacking with respect to the nature of the representation in later stages of the ventral stream supporting object representation, including human object-responsive occipitotemporal cortex (OTC), and the analogue monkey inferotemporal (IT) cortex. In the past decade, this landscape has dramatically changed with the introduction of goal-optimized deep neural networks (DNNs; Krizhevsky et al., 2012; Yamins et al., 2014). These models have revolutionized our methodological capacity to explore the format underlying these late stage visual representations, and provided new traction for empirically testing the impact of different pressures guiding high-level visual representation formation (Kriegeskorte, 2015; Eickenberg et al., 2017; Kay, 2018; Serre, 2019; Richards et al., 2019; Cao and Yamins, 2021; Doerig et al., 2023; Kanwisher et al., 2023).

Landmark findings have demonstrated that deep convolutional neural networks – trained on a rich natural image diet, with the task of object categorization – learn features that predict neural tuning along the ventral visual stream (Yamins et al., 2014; Kriegeskorte, 2015; Güçlü and van Gerven, 2015; Yamins and DiCarlo, 2016; Eickenberg et al., 2017). For example, the responses of single neurons in monkey IT cortex to different natural images can be captured by weighted combinations of internal units in DNNs with greater accuracy than handcrafted features (Yamins et al., 2014). The predictive capacity of these single-neuron encoding models has been further validated in experiments that use these models to synthesize visual stimuli capable of driving neural activity beyond the range evoked by handpicked natural images (Bashivan et al., 2019; Xiao and Kreiman, 2020). The same encoding model procedures carried out using functional magnetic resonance imaging (fMRI) data in humans have shown similarly strong voxel-wise encoding and population-level geometry modeling (Khaligh-Razavi and Kriegeskorte, 2014; Güçlü and van Gerven, 2015; Cichy et al., 2016; Eickenberg et al., 2017; Long et al., 2018; Wen et al., 2018; St-Yves and Naselaris, 2018; Storrs et al., 2021), providing further evidence of the emergent correspondence between the structure of biological visual system responses and the internals of visual DNN models.

However, where there was once a paucity of performant, image-computable visual representations to study, there is now an overabundance. New models with increasingly powerful visual representational competencies, and with variable architectures, objectives, image diets, and learning parameters, are now produced almost weekly. More often than not, these models are designed to optimize performance on canonical computer vision tasks, typically with no reference to brain function nor intent to directly reverse engineer brain mechanisms, per se. This has changed the nature of the problem faced by computational neuroscientists trying to understand high-level visual representation, raising new questions for how to proceed. For example, if these DNN models are to be considered direct models of the brain, is there one model neuroscientists should be using until a better one comes along? Or, might there be another way to leverage the model diversity itself for insight into how more general inductive biases, shared among sets of models, lead to more or less brain-like representation?

Neural benchmarking platforms such as Brain-Score, Algonauts, and Sensorium, directly operationalize the research effort to find the most brain-like model of biological vision, and do so with impressive scale and generality (Schrimpf et al., 2018a, 2020; Cichy et al., 2019; Willeke et al., 2022). Crowd-sourcing across neuroscientists and applied machine learning researchers alike, these platforms collect many brain datasets that sample responses from multiple visual areas to a variety of stimuli, allowing users to upload and enter any candidate model for scoring. An automated pipeline fits unit-wise encoding models (under prespecified linking assumptions) to each candidate DNN, and computes an aggregated score across all probe datasets. These neural benchmarking endeavors seem aimed predominantly at identifying the single best predictive model of the target neural system (e.g. mouse primary visual area, primate ventral stream), a goal that that is reflected in the leaderboards of top-ranking models that have become the standard outputs of the pipeline.

Here, however, we take an alternative approach, instead aiming to leverage the diversity and sheer quantity of open-source DNNs to explore broader claims about pressures guiding visual representation formation (Kanwisher et al., 2023), an approach recently termed the “neuroconnecionist research programme” (Doerig et al., 2023). That is, we conceptualize these models *less* as in silico models of the brain with one-to-one correspondence to different regions, and *more* as abstracted visual representation learners, with representational signatures that are either more or less akin to the biological visual system. With this approach, we take as a premise that different DNNs can operationalize different high-level visual formats, where the pressures guiding the format of the representation are jointly shaped by the inductive biases inherent to architecture, task objective, and the visual input itself. Within this framework, training sets of models that vary only in one of these factors, while controlling all others, can be thought of as controlled rearing experiments (Wood et al., 2020), operationalizing different artificial visual systems to explore targeted questions about which variations give rise to a more or less emergent brain-like representation.

In the current work, we harness sources of controlled variation *already present* among pre-existing, open-source models. These opportunistic experiments let us ask broader questions about different representational constraints: For instance, about meso-scale architectural motifs: do convolutional or transformer encoders learn features that better capture high-level visual responses to natural images, holding task and visual experience constant? Or, about the goal of high-level vision: when visual representations are aligned with language representations, does this provide a better fit than purely visual self-supervised objectives, controlling for architecture and visual experience? Or, about visual experience: do certain training datasets (such as faces alone or places alone) lead to more brain-like representational geometry than others? We supplement these experiments with additional analyses that test the explanatory power of more general factors (e.g. effective dimensionality) that have been proposed as underlying increased brain-predictivity, but that do not fit as neatly into the framework of inductive bias. Broadly, the goals of this work are to reveal the relationships between model variations and emergent brain-like visual representation, to provide insight into the principles shaping high-level visual representation in both biological and artificial visual systems, and to articulate the next steps for the neuroconnectionist enterprise, and similar research efforts.

## Results

Our approach involves first searching through pre-trained model repositories and curating distinct *sets* of models that have performant visual capacities, and which provide meaningful controlled variation in key inductive biases such as architecture, task objective, and visual diet (i.e. training dataset). Each of these analyses involves insolating models that vary along only one of these dimensions, while holding the others constant. In total, we examine the degree to which the representations of 224 distinct DNNs can predict the responses to natural images across human occipitotemporal cortex (OTC) in the 7T Natural Scenes Dataset (NSD; Allen et al., 2022), using two different model-to-brain linking methods. Details on all aspects of our procedure are available in the Methods Section.

The full set of all included models and their most relevant metadata is described in Supplementary Information Table 1. Trained models (N = 160) were sourced from a variety of online repositories (Figure 1B). We also collected the randomly-initialized variants (N = 64) of all ImageNet-1K-trained architectures, using the default random initialization procedure provided by the model repository or original authors. We then grouped these models into controlled comparison sets, to enable ‘opportunistic experiments’ between models that differ in only one inductive bias, holding the others constant.

**Table 1:**
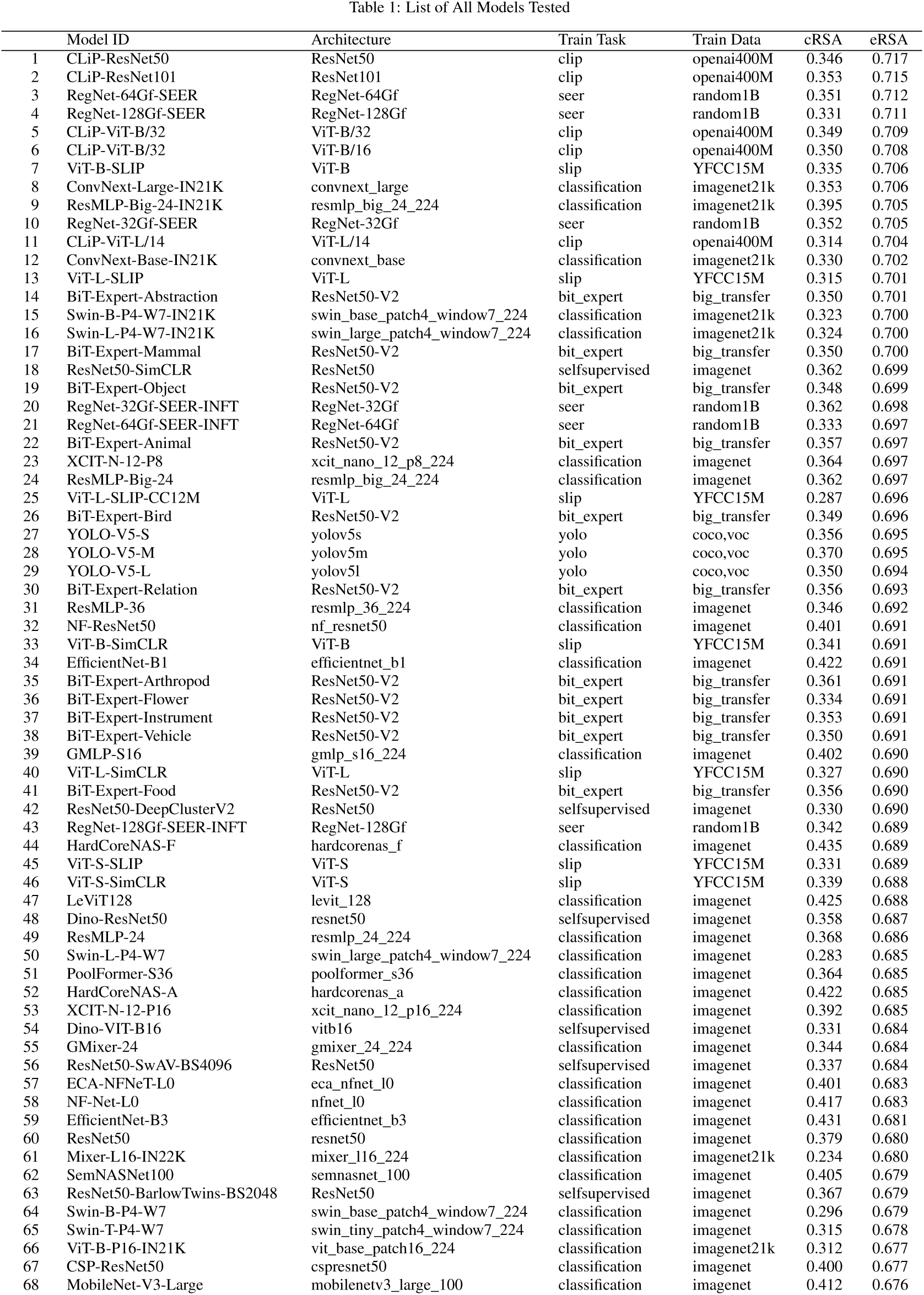
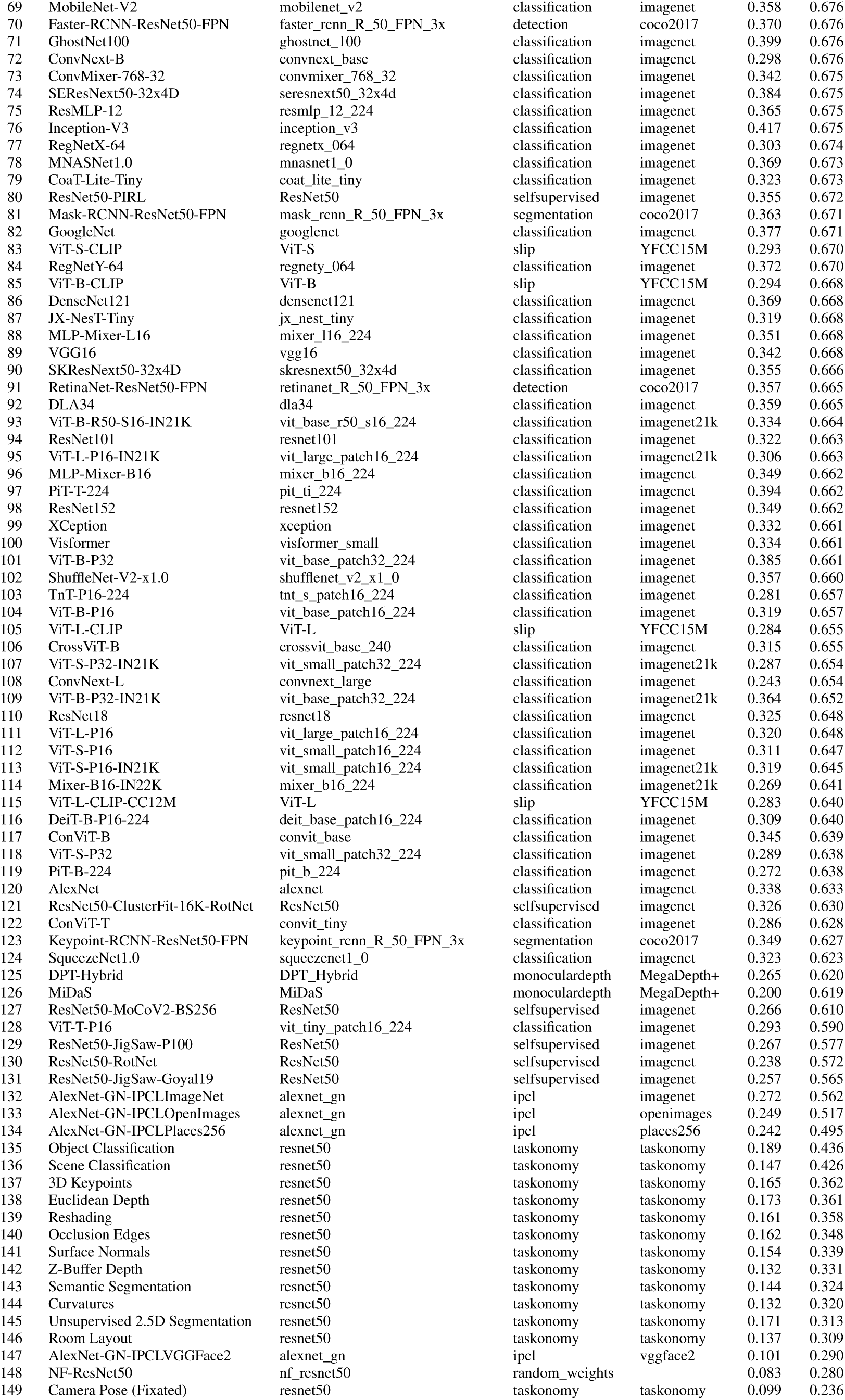
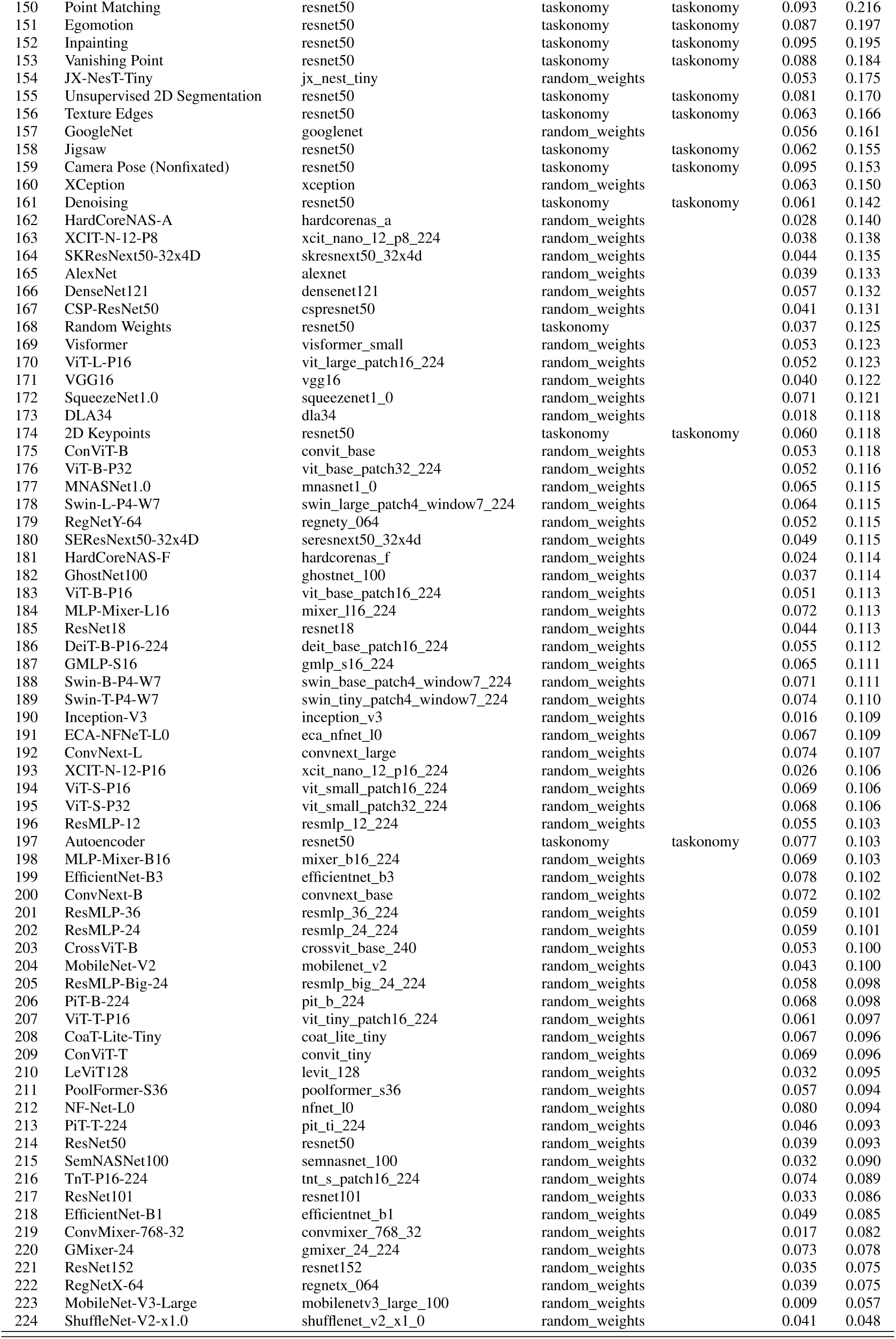
List of All Models Tested

**Figure 1:**
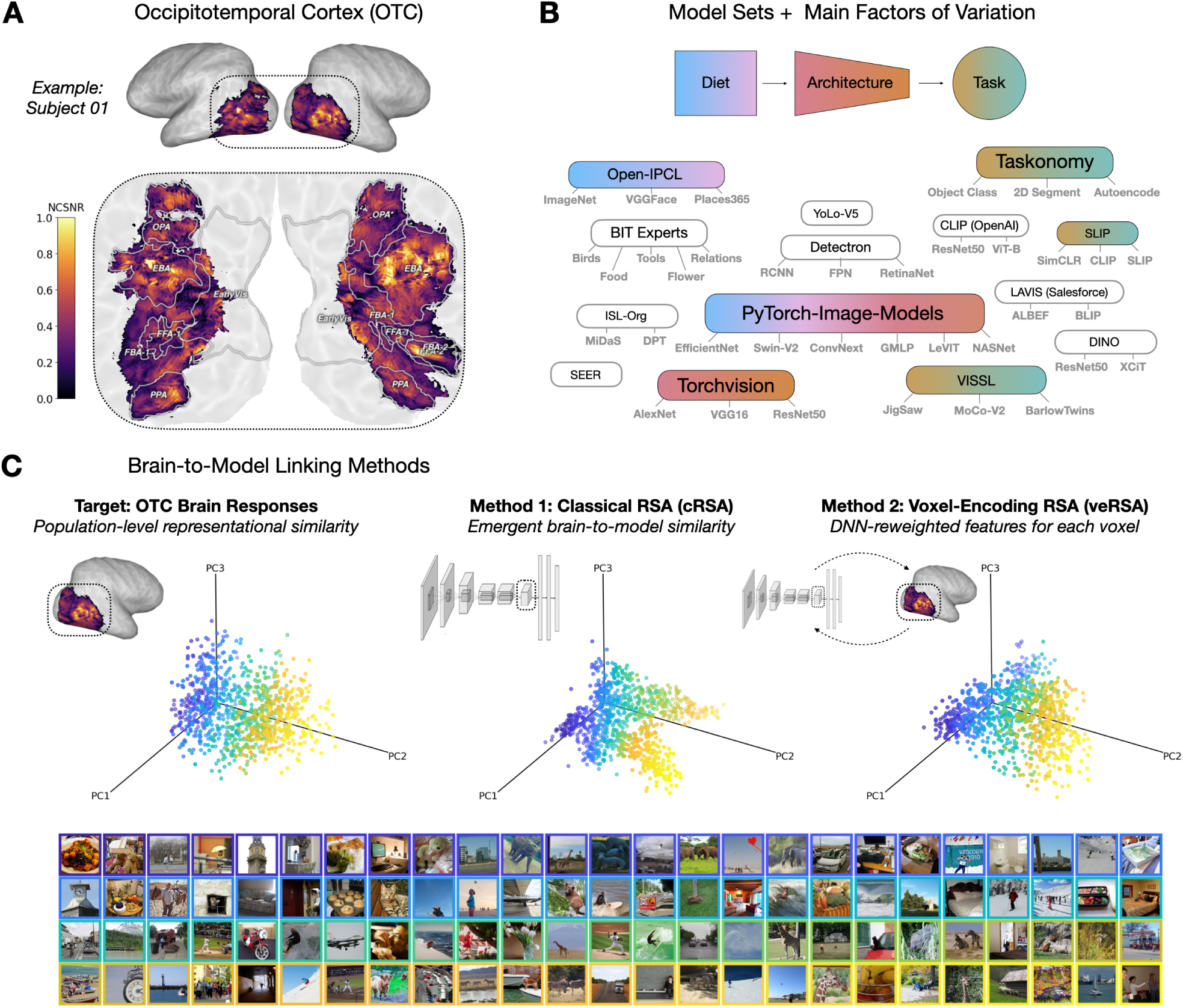
Overview of our approach. ***A*** *The brain region of focus is occipitotemporal cortex (OTC), here shown for an example subject. The voxel-wise noise-ceiling signal-to-noise ratio (NCSNR) is indicated in color. **B** A large set of models were gathered, schematized here by repository, and colored by the main experiments to which they contribute. **C** Brain-linking methods. The left plot depicts the target representational geometry of OTC for 1000 COCO images, plotted along the first three principal components of the voxel space. Each dot reflects the encoding of a natural image, a subset of which are depicted below in a corresponding color outline. The middle panel shows a DNN representational geometry (here the final embedding of a CLIP-ResNet50), plotted along its top 3 principal components. Classical RSA involves directly estimating the emergent similarity between the brain target and the model layer representational geometries. The right plot shows the same DNN layer representation, but after the voxel-wise encoding procedure (veRSA), which involves first re-weighting the DNN features to maximize voxel-wise encoding accuracy, and then estimating the similarity between the target voxel representations and the model-predicted voxel representations*.

The main brain target of our analyses in this work were OTC responses to 1000 natural images in voxels sampled from 4 subjects in the NSD (Figure 1A). We define the OTC sector individually in each participant using a combination of reliability-based (SNR) and functional metrics. Our key outcome measure was the representational geometry of the OTC brain region, captured by the set of 124,750 pairwise distances between 500 test images. The representational geometry in this dataset was highly reliable (noise ceiling mean across subjects: *r_Pearson_* = 0.8, range [0.74, 0.85] across subjects), providing a strong target for arbitrating the relative predictivities of our surveyed models.

We considered two different linking methods to relate the representations learned in models to the response structure measured in brains. Both of these methods predict each individual subject’s OTC representational geometry, using representational similarity analysis (RSA) (Kriegeskorte et al., 2008a). The first method, classical RSA (cRSA), is the more strict linking hypothesis, estimating the degree of correspondence between the brain’s population geometry and the best-fitting model layer’s population geometry, without any feature re-weighting procedures. This measure probes for a fully emergent correspondence between model and brain, making the clear (and reasonable) assumption that as a whole, all units of the DNN layer must contribute equally to capture the population level geometry.

The second method, voxel-encoding RSA (veRSA), tests for the same correspondence, but allows for guided re-weighting of the DNN features (Khaligh-Razavi et al., 2017a; Kaniuth and Hebart, 2022; Konkle and Alvarez, 2022). This method first makes the (also reasonable) assumption that different voxels are likely tuned to different features, and thus, that each voxel’s response profile should be modeled as a weighted combination of the units in a layer, using independent brain data for fitting the encoding model. After fitting, each voxel-wise encoding model is used to predict responses to the test images, for which we compare the predicted population geometry to the observed population geometry of neural responses. Thus, these two mapping procedures provide two distinct measures of the degree to which each model’s internal representations are able to predict population geometry of OTC, with either more strict or more flexible linking assumptions.

### Architecture Comparison

While differences in architecture across models can be operationalized in many different ways (total number of parameters, total number of layers, average width of layers, et cetera), here we focus on a distinct meso-scale architectural motif: the presence (or absence) of an explicit convolutional bias, which is present in convolutional neural networks (CNNs) but absent in vision transformers.

CNN architectures applied to image processing were arguably the central drivers of the last decade’s reinvigorated interest in artificial intelligence (Krizhevsky et al., 2012; LeCun et al., 2015). CNNs are considered to be naturally optimized for visual processing applications that benefit from a sliding window (weight sharing) strategy, applying learned local features over an entire image. These models are known for their efficiency, stability, and shift equivariance (Liu et al., 2022; McGreivy and Hakim, 2022). Transformers, originally developed for natural language processing, have since emerged as major challengers to the centrality of CNNs in AI research, even in vision. Transformers operate over patched image inputs, using multihead attention modules instead of convolutions (Raghu et al., 2021; Naseer et al., 2021).They are designed to better capture long-range dependencies, and are considered in some cases to be a more powerful alternative to CNNs precisely for this reason (Zhou et al., 2021). While many variants of CNNs and some kinds of transformers have appeared in benchmarking competitions (Schrimpf et al., 2020; Cichy et al., 2019; Willeke et al., 2022), comparisons between these models have been largely uncontrolled. Which of these models’ starkly divergent architectural inductive biases lead to learned representations that are more predictive of human ventral visual system responses, controlling for visual input diet and task?

We compared the brain-predictivity scores of 34 CNNs against 21 transformers. Critically, all of these models were trained with the same dataset (ImageNet1K) and the same task objective (1000-way image classification). The results are shown in Figure 2. Surprisingly, we find that both the CNN and transformer architectures account for the structure of OTC responses almost equally well: in the veRSA comparison, the brain predictivity on average was *r_Pearson_* = 0.67 [0.67, 0.68] for convolutional models and *r_Pearson_* = 0.66 [0.65, 0.67] for transformer models. We did find, however, that the aggregate differences between these architectures, while small, were statistically significant (Wald *t*-distribution statistics, see Methods). Specifically, the transformers were on average less predictive than the CNNs, in both the classical and voxel-encoding RSA metrics (cRSA: *β* = -0.04 [-0.05, -0.03], *p* < 0.001; veRSA: *β* = -0.01 [-0.02, -0.00] *p* < 0.001). Thus, the CNNs as a class may introduce an inductive bias that leads to a *slightly* more brain-aligned late-stage visual representation, on average – holding task and visual input diet constant. However, note that the prediction ranges among the surveyed CNN and transformer models were substantially overlapping, so this statistical effect should not be interpreted as a categorical claim that all convolutional models have greater emergent brain predictivity than all transformer models.

**Figure 2:**
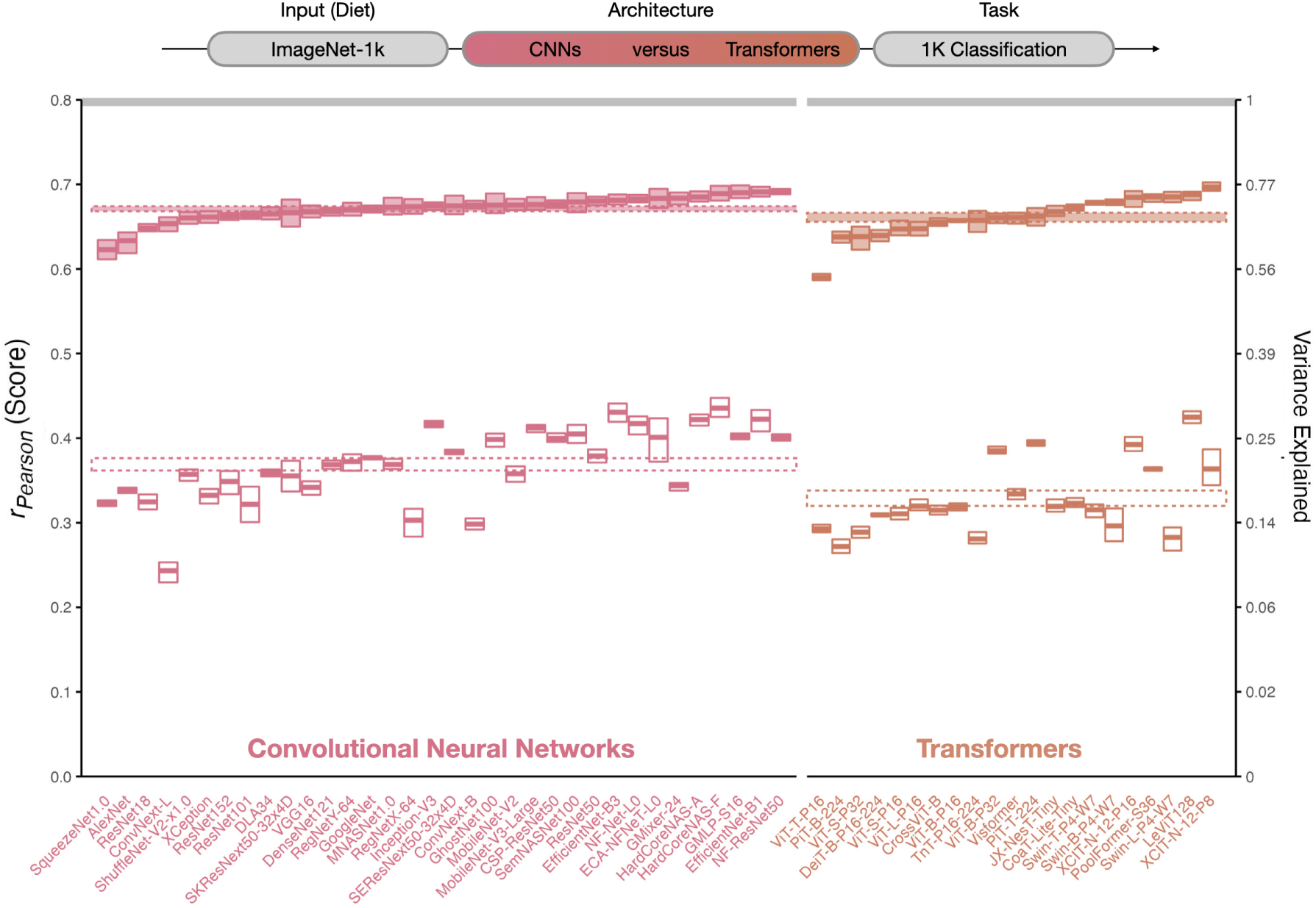
Architecture Variation. *Degree of brain predictivity (r_Pearson_) is plotted for the controlled set of convolutional neural networks (CNNs) and transformer models in our survey. Each small box corresponds an individual model. The horizontal midline of each box indicates the model’s mean score across the 4 subjects, with the height of the box indicating the grand-mean-centered 95% bootstrapped confidence intervals (CIs) (Morey et al., 2008) of the model’s score across subjects. The cRSA score is plotted in open boxes, and the veRSA score is plotted in filled boxes. For each class of model architecture (convolutional, transformer) the class mean is plotted as a striped horizontal ribbon. The width of this ribbon reflects the 95% grand-mean-centered bootstrapped 95% CIs over the mean score for all models in a given set. The aggregate noise ceiling of the occipitotemporal brain data is plotted in the gray horizontal ribbon at the top of the plot, and reflects the mean and 95% CIs of the noise ceilings computed for each individual subject. The secondary y-axis shows explainable variance explained (the squared model score, divided by the squared noise ceiling)*.

Given the dramatic differences between these architectural encoders, we found it surprising how similarly they predicted the structure of the brain responses in high-level visual cortex, which suggests that these models are converging on the same representational format. Worth noting, though, is that the cRSA brain-predictivity scores were both much lower and more variable than the veRSA scores. This implies that feature re-weighting is playing a substantial role in the degree to which these models capture the representational geometry of the high-level visual system. This observation is also consistent with the possibility that the learned representations of these models all capture similar representational sub-spaces after feature re-weighting. We return to this possibility analytically in the “Model-to-Model Comparison” Section.

### Task Variation

Next, we examined the impact of different task objectives on the emergent capacity of a model to predict the similarity structure of brain responses. Our case studies here probe the effect of different canonical computer vision tasks (Zamir et al., 2018), the effect of different self-supervised algorithms (Goyal et al., 2021), and the effect of visual representation learning with or without linguistic alignment (Mu et al., 2021). The results of these experiments are summarized in Figure 3.

**Figure 3:**
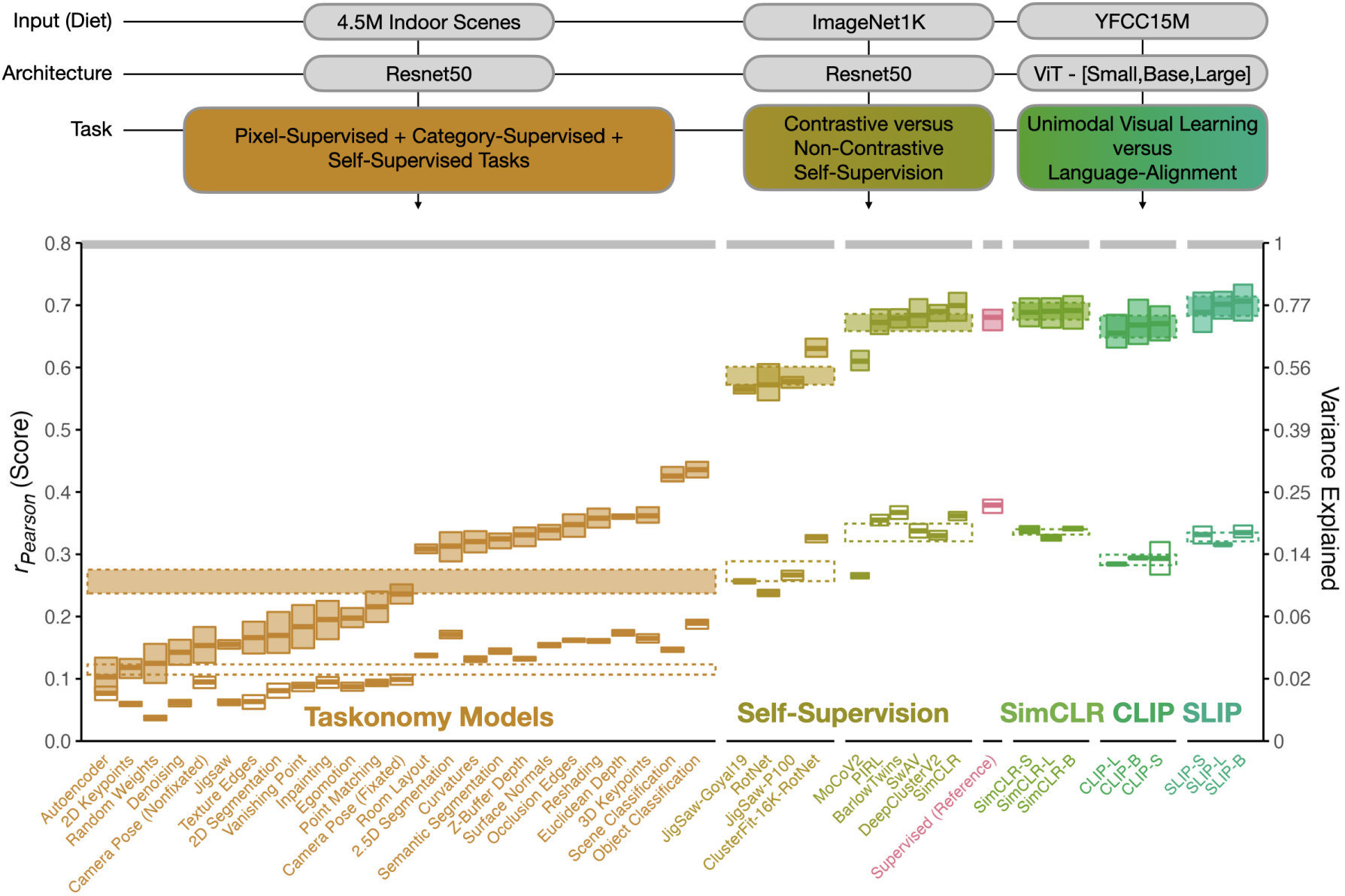
Task Variation. *Degree of brain predictivity (r_Pearson_) is plotted for the sets of models with controlled variation in task. The first set of models shows scores across the ResNet50 encoders from Taskonomy, trained on a custom dataset of 4.5 million indoor scenes. The second set of models shows the difference between contrastive and non-contrastive self-supervised learning ResNet50 models (with a category-supervised ResNet50 for reference), trained on ImageNet1K. The third set of models shows the scores across the vision-only and vision-language contrastive learning ViT-[Small,Base,Large] models from FaceBook’s SLIP Project, trained on the images (or image-text pairs) of YFCC15M. Each small box corresponds an individual model. The horizontal midline of each box indicates the model’s mean score across the 4 subjects, with the height of the box indicating the grand-mean-centered 95% bootstrapped confidence intervals (CIs) of the model’s score across subjects. The cRSA score is plotted in open boxes, and the veRSA score is plotted in filled boxes. The class mean for each distinct set of models is plotted in striped horizontal ribbons across the individual models. The width of this ribbon reflects the 95% grand-mean-centered bootstrapped 95% CIs over the mean score for all models in this set. The aggregate noise ceiling of the occipitotemporal brain data is plotted in the gray horizontal ribbon at the top of the plot, and reflects the mean and 95% CIs of the noise ceilings computed for each individual subject. The secondary y-axis shows explainable variance explained (the squared model score, divided by the squared noise ceiling)*.

### The Taskonomy Models

First, we examined the Taskonomy models (Zamir et al., 2018; Sax et al., 2019) – an early example of controlled model rearing. The Taskonomy models were originally designed to test how well learned representations trained with one task objective transfer to other tasks. Each of these 24 models were trained on different tasks spanning a range of unsupervised and supervised objectives (e.g. autoencoding, depth prediction, scene classification, surface normals, edge detection), some requiring pixel-level labeling and others requiring a single label for the whole image. In all cases, the base encoder architecture is a ResNet50, modified with a specialized projection head to fit the task-specific output. In this analysis, we consider only the feature spaces of the base ResNet50 encoders. The dataset on which the Taskonomy models are trained is a large dataset in terms of raw images, consisting of 4.5 million images, but depicts only images of indoor scenes, with any images of people excluded.

Comparing the brain-predictivity scores of the Taskonomy models, we make two key observations. The first is that across the different task objectives, there was indeed a large range of scores. The least brain-aligned task – autoencoding – yielded a *r_Pearson_* = 0.077 [0.066, 0.085] in cRSA and *r_P earson_* = 0.103 [0.096, 0.11] in veRSA, while the most brain-aligned task – object classification–yielded a *r_P earson_* = 0.189 [0.178, 0.201] in cRSA and *r_Pearson_*= 0.436 [0.419, 0.454] in veRSA).

The second key observation is that the overall range of brain-predictivity scores among these models was relatively low – even for the highest-scoring tasks: for object classification, the veRSA score was only *r_Pearson_* = 0.44 [0.42, 0.45]. For reference, a standard ResNet50 architecture also trained on image classification, but over the ImageNet dataset, shows an average brain predictivity of *r_Pearson_*= 0.68 [0.63, 0.72] (*t_Student_*(3) = -14.3, p < 0.001). Note that this difference manifests *in spite* of Taskonomy’s larger training set (*∼*4.5M images), nearly thrice that of the ImageNet1K (*∼*1.2M images). This observation leads to the hypothesis that the relatively weaker overall brain-predictivity scores for Taskonomy models is related to insufficient diversity of the Taskonomy images^2^.

### Self-Supervised Algorithms

Early variants of self-supervised objectives involved learning representations by predicting image rotations (RotNet) or unscrambling images (JigSaw). More modern variants of self-supervised objectives operate by learning to represent individual images distinctly from one another in an embedding space (SimCLR, BarlowTwins). In particular, contrastive learning objectives typically build a high-level embedding of images by representing samples (augmentations) of the same image nearby in feature space, and far from the representations of other images. Critically, when these learned representations are probed on the canonical computer vision task of image classification, emergent categorization capacity is nearly comparable to that of models trained with category supervision (Geirhos et al., 2020; Chen et al., 2020). Additionally, these contrastive learning models have also been shown to predict brain activity on par with category-supervised models in mice, humans, and non-human primates (Conwell et al., 2021; Nayebi et al., 2021; Zhuang et al., 2021; Konkle and Alvarez, 2022). However, there has not been a systematic comparison of brain predictivity for models trained with these different kinds of self-supervised objectives.

In this experiment, we examine the different brain predictivities of contrastive versus non-constrastive self-supervised learning methods using a suite of 10 models from the VISSL model zoo (Goyal et al., 2021). Each of these models are trained using a different method of self-supervision, but all with a ResNet50 architectural backbone, and a training dataset that consists of the images (but not the labels) of the ImageNet1K dataset. We divide the models of this set into two groups: models that employ instance-level contrastive learning (N = 6: PIRL, DeepClusterV2, MoCoV2, SwaV, SimCLR, BarlowTwins) and models that do not (N = 4: RotNet, two Jigsaw variants, ClusterFit).

These two different classes of self-supervised learning objectives yield significantly different brain predictivities. Instance-level contrastive learning objectives lead to more brain-predictive representations than non-contrastive objectives by a significant, midsize margin (cRSA: *β* = -0.06 [-0.04, -0.09], *p* < 0.001; veRSA: *β* = -0.09 [-0.07, -0.11] *p* < 0.001). And, consistent with prior work (Zhuang et al., 2021; Konkle and Alvarez, 2022), most instance-level contrastive objectives provide comparable brain predictivity to the matched category-supervised model, holding architecture and visual input diet constant. (For example, the average predictivity of VISSL’s ResNet50-BarlowTwins was 0.367 [0.353, 0.381] in cRSA and 0.679 [0.637, 0.720] in veRSA; the predictivity of Torchvision’s ImageNet1K-trained category-supervised ResNet50 was *r_Pearson_* = 0.379 [0.363, 0.39] in cRSA and 0.680 [0.640, 0.718] in veRSA. These models share identical architectures in PyTorch.) Broadly, these results highlight the potential for understanding biological visual representation through deeper exploration of instance-level contrastive objectives, which focus on learning invariances over samples from the same image, while also learning features that discriminate distinct individual images.

### Language Alignment (The SLIP Models)

Another recent development in self-supervised contrastive learning involves leveraging the structure of another modality – language – to influence representations learned in vision. The preeminent example of this ‘language-aligned’ learning is OpenAI’s CLIP model (Contrastive Language-Image-Pretraining Radford et al., 2021), which builds representations designed to maximize the cosine similarity between the latent representation of an image and the latent representation of that image’s caption. In computer vision research, CLIP has demonstrated remarkable zero-shot generalization to image classification task variants without further retraining. Indeed, across all the models in our set, the model with the highest brain-predictivity is OpenAI’s CLIP-ResNet50 model, with 4 other CLIP-trained models from OpenAI also in the top 10 (see Supplementary Information SI.1). If the constraint of language alignment is key in building more brain-aligned representations, this constitutes a development of deep theoretical import when considering the pressures guiding late stage representations in the ventral visual hierarchy.

A critical issue in using OpenAI’s CLIP model for brain prediction, however, is the fact that OpenAI not only introduced a new objective (image-text alignment via contrastive learning), but also a massive new training dataset (a highly curated and proprietary dataset of 400 million image-text pairs, which has yet to be made available for public research). This makes the direct comparison of CLIP to other models empirically dubious, as it leaves open whether gains in predictivity are attributable to language alignment per se, to massive dataset differences, or the interaction of the two.

Fortunately, Meta AI has since released a set of models – the SLIP models (Mu et al., 2021) – that compare self-supervised contrastive learning models with and without language alignment, controlling for training dataset and architecture. These models are trained with one of 3 learning objectives: pure self-supervision (SimCLR), pure language alignment (CLIP), or a combination of self-supervision and language alignment (SLIP). All models consist of a Vision Transformer backbone (three sizes: Small [ViT-S], Base [ViT-B], & Large [ViT-L]), and are all trained on the YFCC15M dataset (15 million images or image-text pairs). Thus, any differences in brain predictivity across these models reflect the impact of language alignment per se, holding image diet and architecture constant.

Comparing OTC predictivity across these models (Figure 3: SimCLR, CLIP, SLIP), we find that all models have relatively high brain-predictivity scores, with no substantial differences between objectives with and without language alignment. If anything, we see a slight, but significant *decrease* in accuracy for the pure language-aligned CLIP objective. Specifically, in a linear model regressing brain predictivity on the interaction of model size (ViT-[S,B,L]) and task (SimCLR, CLIP, SLIP) plus an additive effect of subject ID, the only significant experimental effect is a slight decrease in the predictive accuracy of the pure CLIP model relative to SimCLR in both metrics (cRSA: *β* = -0.05 [-0.06, -0.03], *p* < 0.001; veRSA: *β* = -0.02 [-0.04„ -0.01] *p* = 0.005). We find no effect of the model size, nor an interaction with the task objective.

These results thus lead to the somewhat surprising conclusion that, in terms of brain predictivity, the superior performance of OpenAI’s CLIP models is likely due to the expanded and proprietary image database, rather than the influence of language alignment per se (though it is possible the benefits of language alignment only emerge at datasets of this more massive scale). Likewise, emerging insights in the machine learning community are also finding the visual representational robustness of OpenAI’s CLIP is almost exclusively conferred by image dataset (Wortsman et al., 2022). Our conclusions about the negligible impact of language alignment on brain predictivity also directly contrast with those of recent papers that conclude the opposite, based on uncontrolled comparisons of CLIP-ResNet50 against ImageNet-1K-trained ResNet50 (Wang et al., 2022). Thus, our work also strongly underscores the need for more empirically controlled model sets (like those of SLIP) that could better arbitrate questions about the emergent properties of language-aligned models trained on massive datasets.

### Input Variation

We next directly examined the impact of a model’s input diet on brain predictivity. Here, we define a model’s input diet as the images used to train the model, irrespective of whether their labels factor into the training procedure. While there are many different datasets used across the 160 models in our model set, there were actually relatively few subsets that enabled controlled comparison of the impact of dataset while holding architecture and objective constant. In the two controlled experiments possible, we examined the relative brain predictivity of architecture-matched models trained on ImageNet1K versus ImageNet21K, and of instance-level contrastive learning models trained on datasets of faces, places, objects. The results of these experiments are summarized in Figure 4.

**Figure 4:**
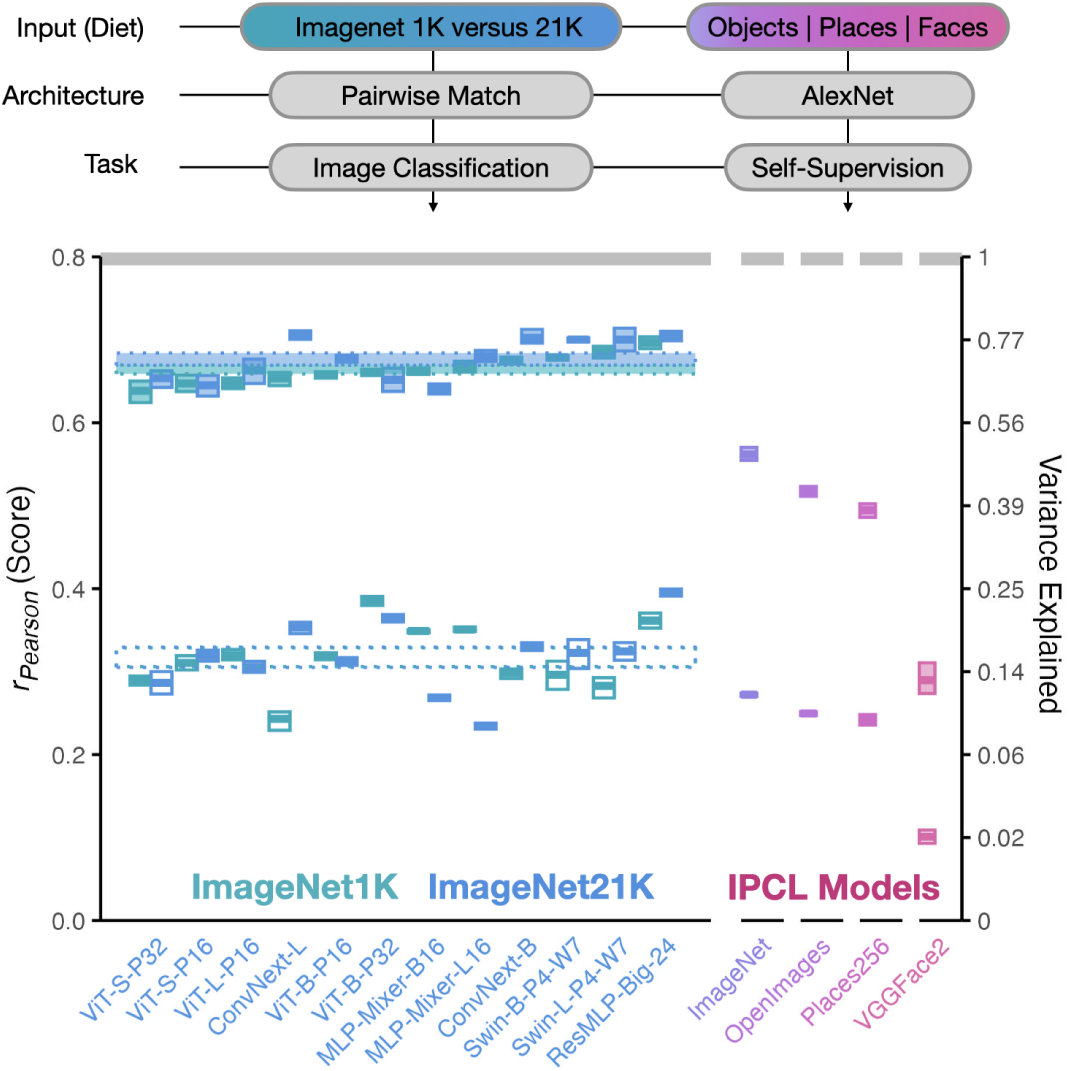
Input Variation. *Degree of brain predictivity (r_Pearson_) is plotted for the sets of models with controlled variation in input diet. The first set of models shows scores across paired model architectures trained either on ImageNet1K or ImageNet21K (a ∼13x increase in number of training images). The second set of models shows scores across 4 variants of a self-supervised IPCL-AlexNet model trained on different image datasets. Each small box corresponds an individual model. The horizontal midline of each box indicates the model’s mean score across the 4 subjects, with the height of the box indicating the grand-mean-centered 95% bootstrapped confidence intervals (CIs) of the model’s score across subjects. The cRSA score is plotted in open boxes, and the veRSA score is plotted in filled boxes. The class mean for each distinct set of models is plotted in striped horizontal ribbons across the individual models. The width of this ribbon reflects the 95% grand-mean-centered bootstrapped 95% CIs over the mean score for all models in this set. The aggregate noise ceiling of the occipitotemporal brain data is plotted in the gray horizontal ribbon at the top of the plot, and reflects the mean and 95% CIs of the noise ceilings computed for each individual subject. The secondary y-axis shows explainable variance explained (the squared model score, divided by the squared noise ceiling)*.

### Imagenet1K versus Imagenet21K

The first controlled dataset comparison we performed was between models trained either on ImageNet1K or ImageNet21K. Deep learning is notoriously data-intensive, and most if not all DNNs are known to benefit from more training samples when it comes to overall classification accuracy (Kaplan et al., 2020; Goyal et al., 2022). ImageNet21K is a dataset of *∼*14.2 million images across 21,843 classes (many of which are hierarchically labeled). The more popular version of ImageNet1K (with 1000 classes) is a *∼*1.2 million image subset of this larger dataset. ImageNet21K is considered by some to be a more diverse dataset (Puigcerver et al., 2020; Ridnik et al., 2021), though exact quantification of this diversity has proven difficult. Does training on this larger and purportedly more diverse dataset lead to increased brain predictivity?

We considered 14 pairs of models from the PyTorch-Image-Models (Wightman, 2019) repository with matched architecture and task-objective, trained either on the ImageNet1K or the ImageNet21K dataset. Performing paired statistical comparisons between their prediction levels, we find no effect of input diet on resulting brain predictivity, in either metric (cRSA *β* = 0.0 [-0.03, 0.03], p = 0.957; veRSA *β* = 0.01 [0.00, 0.03], p = 0.147). This result highlights first that the raw quantity of training images does not necessarily lead to increased brain-predictivity. This result also highlights that whatever the increased diversity of ImageNet21K, it does not manifest in this particular comparison to high-level visual representations in human OTC.

### Objects versus Places versus Faces

The second controlled dataset experiment we perform is based on the set of IPCL-Alexnet models, trained on 4 different datasets: object-focused image sets (ImageNet-1K, OpenImages), scene-focused image sets (Places365), or face-focused image sets (VGGFace2). This model set uses the same architecture and the same self-supervised, instance prototype contrastive learning objective (Konkle and Alvarez, 2022). Notably, these models are trained without labels – further deconfounding task and visual input diet.

In this model set, we find that the ImageNet diet leads to the overall highest brain predictivity: relative to the ImageNet-trained model, the OpenImages-trained model yields a decrease in score of *β* = -0.02 [-0.03,-0.01], p = 0.002 in cRSA, and *β* = -0.04 [-0.07, -0.02], p < 0.001 in veRSA). The Places-365 trained model yields a decrease in score of *β* = -0.03 [-0.04,-0.02], p < 0.001 in cRSA, and *β* = -0.07 [-0.09, -0.04], p < 0.001 in veRSA. Finally, the VGGFace2-trained model is significantly worse than the ImageNet-trained model by a substantial margin in both metrics (cRSA *β* = -0.17 [-0.18,-0.16], p < 0.001; veRSA *β* = -0.27 [-0.30, -0.25], p < 0.001). These results highlight that – at least for this model architecture and objective – the visual diet leads to substantial variation in brain predictivity.

Finally, we note that in each of these cases, the VGGFace2 and Places-365 dataset actually contain more images than ImageNet (*∼*2.75x and *∼*1.5x, respectively), again underscoring that the quantity of images is not the relevant factor here. Broadly, these results hint yet again at the importance of a latent dataset diversity factor that has yet to be quantified. A speculative but intriguing reverse inference suggests that ImageNet, OpenImages, Places-365, and VGGFace2 may be ranked in terms of diversity from greatest to least, based on their ability to capture the representational structure of OTC in response to hundreds of natural images.

### Impact of Training

Research in network initialization techniques have in recent years led to the development of models with randomized, hierarchical representations (effectively, multiscale hierarchical random projections) that are sometimes powerful enough to serve as functional substitutes for trained features in a variety of tasks (Gallicchio and Scardapane, 2020). Past experiments with brain-to-model mappings, for example, have suggested that randomly-initialized models may in certain cases (and for certain species) be as predictive of the brain as trained models (Cadena et al., 2019).

While subsequent analyses have since suggested such findings apply only in a limited set of architectures (Conwell et al., 2021), we did include another such test here: for every trained model architecture we test from the TorchVision and PyTorch-Image-Models repositories, we also test one randomly initialized counterpart (N = 64). This paired statistical test confirms that training has a resounding effect on brain predictivity in both metrics (cRSA *β* = 0.30 [0.29, 0.31], p < 0.001; veRSA *β* = 0.56 [0.55, 0.75], p < 0.001) – the single largest and most robust effect across all of our analyses. In short, training matters.

### Overall Variation across Models

In all analyses to this point, we have focused on understanding variation in brain-predictivity scores through targeted model comparisons, with controlled differences in inductive biases defined by their architecture, task, or input diet. We next focus on understanding variation in brain predictivity across *all* models in our survey.

Across the full set of 224 models we tested (including randomly-initialized variants) and both metrics, we observe predictivity scores that span nearly the full range possible between 0 and the noise ceiling (*r_Pearson_* = 0.8 [0.741, 0.847]). (See Supplementary Information SI.3 for an analysis that considers the stability of these scores across subjects). Figure 5 shows the brain predictivity of all 224 models. From this graph, it is clear that a large number of models perform comparably well, with 126 models yielding veRSA brain-predictivity scores that differ by less than *r_P earson_* = 0.1. (A bootstrapped segmented regression analysis over these scores indicate a break point at rank 124 [123.6, 124.7] /224, corresponding to scores of *r_P earson_* = 0.623 and lower). The ranks beyond this elbow are populated almost entirely by models trained on image diets less diverse than ImageNet (e.g. Taskonomy), and untrained models^3^.

**Figure 5:**
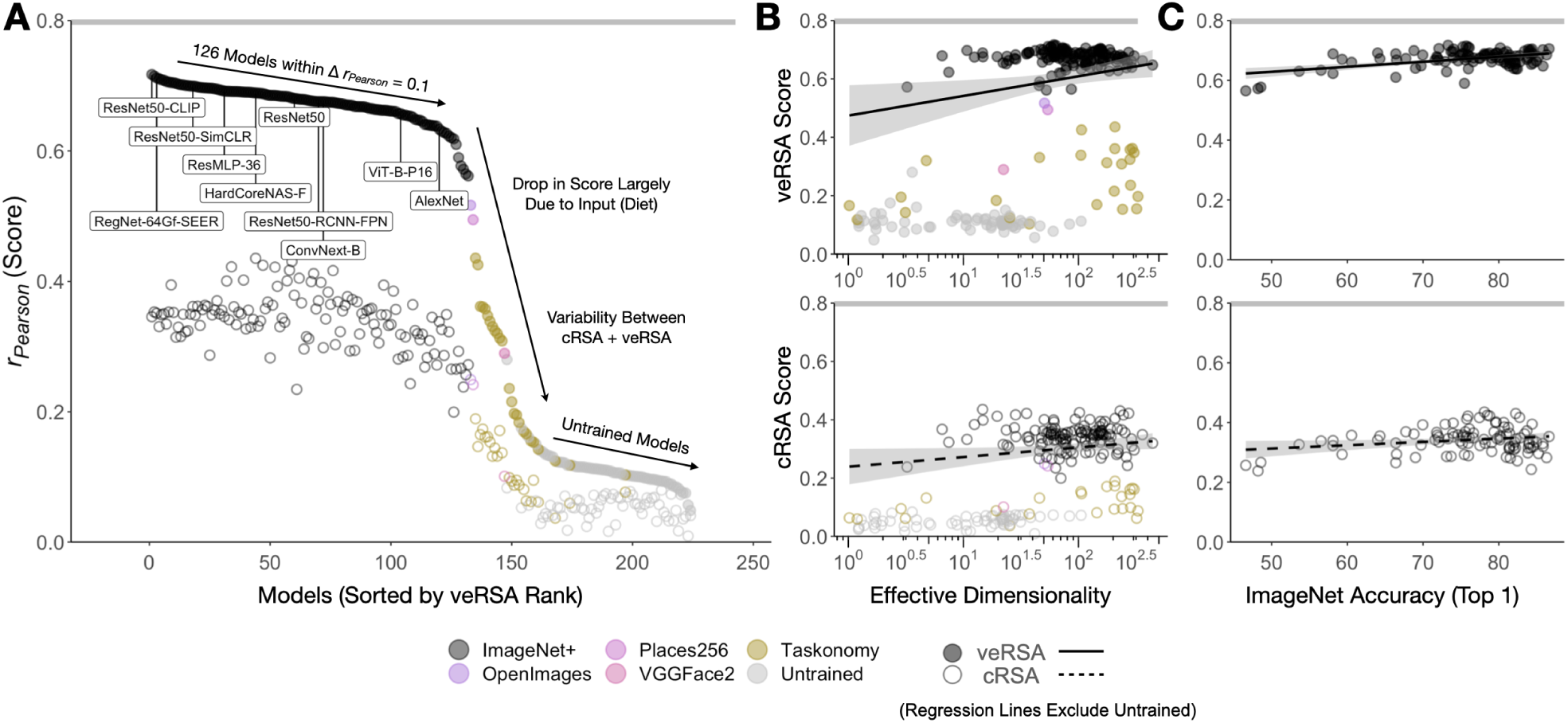
Overall Model Variation. ***A*** *Brain predictivity is plotted for all models in this survey (N = 224), sorted by veRSA score. Each point is the score of a single model, plotted for both cRSA (open) and veRSA (filled) metrics. Models trained on different image sets are labeled in color. **B** Brain predictivity is plotted as a function of the effective dimensionality of the most predictive layer, with veRSA scores in the top panel and cRSA scores in the bottom panel. The regression line is fit only on trained variants of the models (excluding untrained variants). **C** Brain predictivity is plotted as a function of the top-1 ImageNet1K-categorization accuracy for the models (N = 108) whose metadata includes this measure (veRSA, top panel; cRSA bottom panel). The noise ceiling of the OTC brain data is shown as the gray horizontal bar at the top of each plot*.

Given this degree of variation, we next focused on three more general factors that have been hypothesized to account for a model’s emergent brain predictivity beyond differences in inductive bias: effective dimensionality, classification accuracy, and parameter count.

### Effective Dimensionality

Recent work in DNN modeling of high-level visual cortical responses in both human and non-human primates has suggested a general principle, wherein model representations with a higher ‘latent’ or ‘effective’ dimensionality (Del Giudice, 2021) are more predictive of high-level visual cortex (Elmoznino and Bonner, 2022). Effective dimensionality (ED), in this case, is a property of manifold geometry defined as the “continuous measurement of the number of principal components needed to explain most of the variance in a dataset” (Elmoznino and Bonner, 2022).

In this analysis, we sought to test whether the relationship between ED and brain predictivity holds across the full set of models in our survey. To do so, we computed the ED of the most OTC-aligned layer representations from each model (as measured by our veRSA metric), using the same 1000 COCO images from our main analysis (see Methods and Supplementary Information for details). The relationship between model layer effective dimensionality and its corresponding brain predictivity is shown in Figure 5.

We first considered variation in ED across all models – both trained and randomly-initialized, akin to Elmoznino and Bonner (2022). From this perspective, ED appears to be a significant, moderately high predictor of each model’s veRSA score: *r_Spearman_* = 0.489 [0.381, 0.580], *p* < 0.001, across 1000 bootstraps of the sampled models. However, in our data, this relationship seems to be driven almost entirely by the predictivity differences between trained and untrained models. When we computed the relationship between ED and prediction scores for trained and randomly-initialized models separately, variation in the ED of trained models showed no correlation with the veRSA score (*r_Spearman_* = -0.063 [-0.31, 0.099], *p* = 0.692). Similarly, variation in ED among randomly-initialized model layers produced a non-significant, slightly negative correlation with OTC prediction (*r_Spearman_* = -0.142 [-0.307, 0.118], *p* = 0.077). See Supplementary Information SI.2 for a more detailed exploration of the impact of ED in our data in relation to Elmoznino and Bonner (2022), and the different analytical choices that may underlie the divergence in our results.

Overall, our analyses suggest that this particular effective dimensionality metric is not a general principle explaining emergent brain predictivity. This is by no means a rejection of the more general hypothesis that the geometric and statistical properties of neural manifolds might well transcend inductive bias as predictors of brain similarity, but it does suggest that we may need different metrics (e.g. Garrido et al., 2022; Yerxa et al., 2023) to unveil these underlying principles moving forward.

### Classification Accuracy

Early studies of brain-predictive DNNs provided evidence of a link between a model’s ability to accurately perform object categorization and its capacity to predict the responses to neurons along the primate ventral visual stream (Yamins et al., 2014). However, more recent work suggests that across modern neural network architectures, ImageNet top-1 performance acts as a weaker or even null predictor of brain similarity (Schrimpf et al., 2018b, 2020; Konkle and Alvarez, 2022; Linsley et al., 2023). We next examined this relationship in our human OTC data, considering the N = 99 models for which we have the ImageNet-1K top-1 categorization accuracy available. Figure 5 (rightmost panels) plots a model’s top-1 accuracy against the brain predictivity of that model’s best-predicting layer. We find little to no relationship between classification accuracy and brain predictivity across these models, in either cRSA or veRSA (*r_Spearman_* = -0.0057, *p* = 0.96 in cRSA; *r_Spearman_* = 0.17, *p* = 0.088 in veRSA; rank-order correlation is appropriate given the non-normality of top-1 accuracies). Our results also do not show an obvious plateau in accuracies above 70% (c.f. Schrimpf et al., 2018b); rather, the models in our comparison set have a relatively wide range of top-1 accuracies with a relatively restricted range of brain-predictivity scores. However, we note that trained models far outperform untrained models. Thus, task performance likely plays an important role for high-level visual representational quality to some degree, but such effects may no longer be captured by the fine-grained metric of top-1 object recognition accuracy that was once the primary standard of this particular competence.

### Number of Trainable Parameters

Model size is another attribute commonly suspected to influence many DNN outcome metrics, including emergent predictivity of human brain and behavioral data (Kaplan et al., 2020; Geirhos et al., 2021; Schrimpf et al., 2020; Muttenthaler et al., 2022; Dehghani et al., 2023), and is typically measured via quantification of depth, width, or total number of parameters. Here, we test for differences in brain predictivity as a function of total parameter count for all models, excluding the Taskonomy and VISSL ResNet50s, as well as IPCL-AlexNet models, which (sharing the same architectures) do not vary in their total number of parameters. We find a somewhat irregular set of patterns. As parameter count increases in *trained* models, there is a significant, midsize *decrease* in cRSA brain-predictivity (*r_Spearman_* = -0.45, *p* = 0.026*e^−^*^7^, and a non-significant *increase* in veRSA brain-predictivity (*r_Spearman_* = 0.14, *p* = 0.129). As parameter count increases in *untrained* models, there is a significant, small-to-midsize increase in cRSA score (0.28, *p* = 0.0225), coupled with another non-significant increase in veRSA score (0.083, *p* = 0.513). In short, there is no consistent influence of total trainable parameter count on subsequent brain predictivity.

### Model-to-Model Comparison (RSA)

Considered in the aggregate, the opportunistic experiments and overarching statistics in this work are underwritten by over 1.8 billion regression fits and 50,300 representational similarity analyses. Given the scale of our analyses and the surprisingly modest set of significant relationships that arise from them, it would not be entirely unreasonable at this point to arrive at a somewhat deflationary conclusion – that almost none of the factors *hypothesized* as central to the better modeling of brains actually translate to more brain-aligned representations *in practice*. Convolutions, language alignment, effective dimensionality – all concepts of deep theoretical import – make almost no difference when it comes to predicting how well a model will ultimately explain high-level visual cortical representations in the largest such fMRI dataset gathered to date. 126 models – many of which appear to differ fundamentally in their design – have veRSA brain-predictivity scores within *r_P earson_* = 0.1 of a notably high noise ceiling.

A key question for this research enterprise, then, is to understand whether the different architectures, objectives, and diets actually lead to different representations in the first place. For example, are all of the most brain-aligned models converging on effectively the same representational structure?

To explore this question, we performed a direct model-to-model similarity analysis. Specifically, we computed the pairwise similarities of the most brain-aligned feature spaces from each model, and compared each of these model representations to those from all other models. Critically, we did this using two methods. The first method operates over the classical representational similarity matrices (cRSMs) from each model (i.e. the unweighted RSMs), assessing the pairwise similarity of each model’s cRSM to every other model’s cRSM. The second method operates over the voxel-wise encoding RSMs from each model (i.e. the RSM that is produced after feature reweighting), assessing the pairwise similarity of each model’s veRSM to each other model’s veRSM. Taken together, the output of this analysis is effectively two model-to-model meta-RSMs whose constituent pairwise similarities allow us to assess how similar different models’ most brain-aligned layer representations are, with and without being linearly reweighted to predict fMRI responses.

The results are shown in Figure 6. Considering only the top 124 most brain-predictive models, we find that a direct pairwise comparison of their most brain-aligned layers yields substantial variation in representational similarity, with a range that extends from *r_P earson_* = -0.107 to 0.983 (mean = 0.448, SD = 0.148). Thus, these model layers express substantially different representational structure in response to the 500 natural image probe set we use in this analysis. However, the feature-reweighted model representational structure showed a much tighter distribution (mean = 0.881, SD = 0.0313). Thus, the linear reweighting of DNN features in veRSA seems to reveal a remarkably brain-aligned, shared subspace in almost all trained models (or at least, those models trained on a sufficiently diverse image diet).

**Figure 6:**
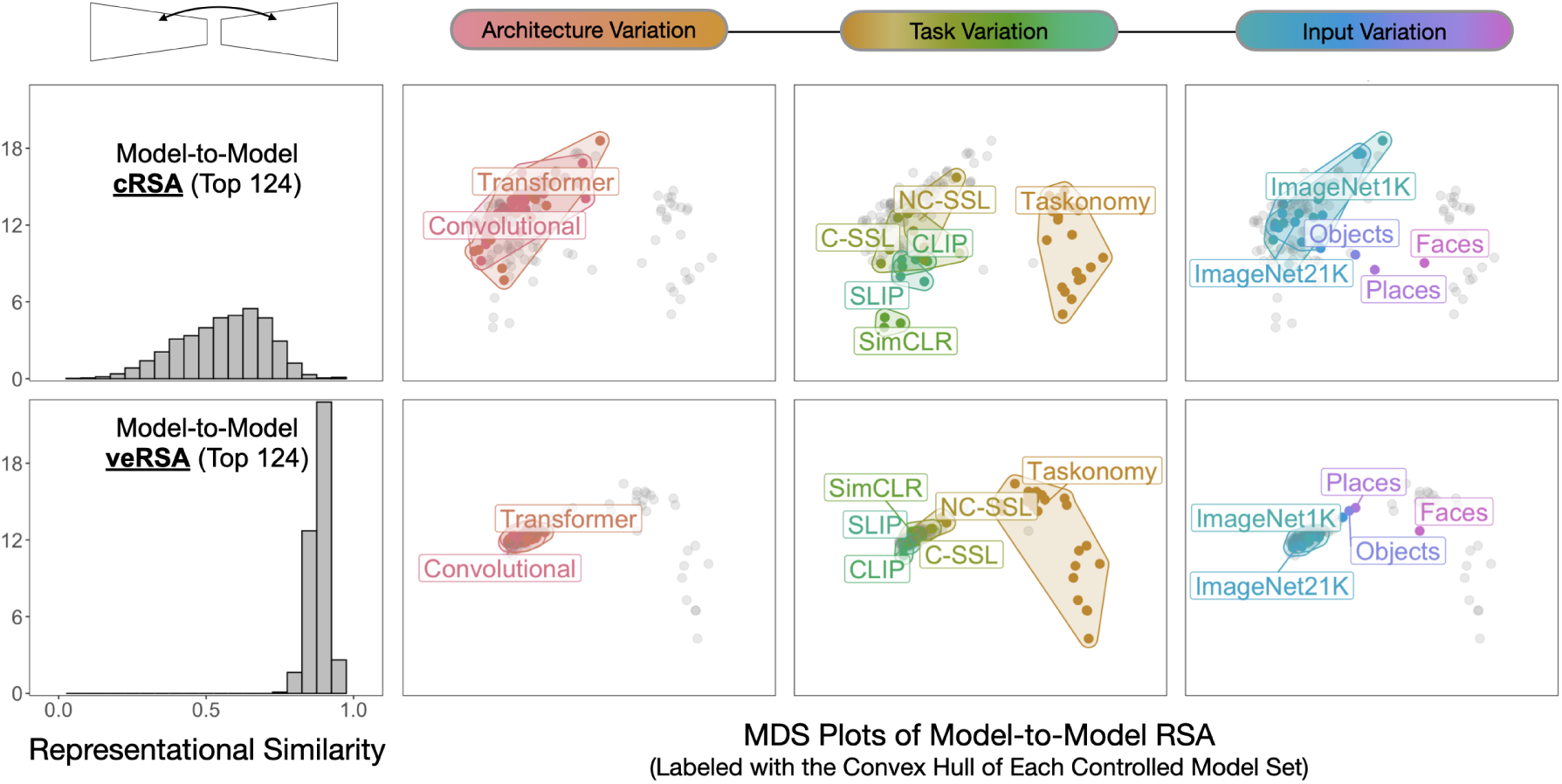
Model-to-Model Comparison. Leftmost Panel: Histogram of the pairwise model-to-model representational similarity for the 124 highest-ranking trained models in our survey. The top panel indicates direct layer-to-layer comparisons, while the bottom panel reflects the feature-reweighted layer-to-layer comparisons. Rightward Panels: Results of a multidimensional scaling (MDS) analysis of the model-to-model comparisons, where model layers with more similar representational structure are plotted more closely to one another. The 3 plots in each row show datapoints output from the same MDS procedure (cRSA, top row; veRSA, bottom row), and the columns show different colored convex hulls that highlight the different model sets from the opportunistic experiments. Note the scale of the MDS plots is the same across all panels. NC-SSL and C-SSL correspond to Non-Contrastive and Contrastive Self-Supervised Learning, respectively. Objects, Faces, and Places correspond to the IPCL models trained on ImageNet1K /OpenImages, Places256, and VGGFace2, respectively.

The adjacent subplots in Figure 6 show a multi-dimensional scaling plot of the model-to-model comparisons (using all N = 160 trained models), with the axis scales held constant across all facets. Model layers with more similar representational structure are displayed nearby. These plots make clear that there is a substantial degree of raw representational variation amongst these models, which is dramatically compressed when these feature spaces are mapped onto responses in human OTC.

These findings suggest that the somewhat hidden factor of *model-to-brain mapping method* (and the linking assumptions inherent to these methods) is at least as consequential, if not more consequential, than differences in inductive bias. Indeed, if we treat our metrics themselves (cRSA versus veRSA) as a factor in the same kind of linear regression model we use for our controlled model comparisons, we find that the difference between the two constitutes one of the most substantial effects on brainpredictivity of any we assess (second only to that of trained versus random weights): *β* = 0.23 [0.21, 0.25], *p* < 0.001 for all models, trained and random; *β* = 0.30 [0.29, 0.31], *p* < 0.001 for trained models only.

## Discussion

As performant image-computable representation learners, with accessible internal parameters, deep neural networks offer tools for directly operationalizing and testing hypotheses about the formation of high-level visual representations in the brain. This is arguably the core tenet of theoretical frameworks proposed in recent years to unify deep learning and experimental neuroscience (e.g., Richards et al., 2019), and encapsulated most strongly in what has been called the “neuroconnectionist research programme” (Doerig et al., 2023). According to these frameworks, controlled manipulation across DNNs that learn with different inductive biases, combined with appropriate linking methods and predictivity metrics, has the potential to unveil the pressures that have shaped the representation we measure in the brain, answering questions about “why” these representations appear as they do (Kanwisher et al., 2023).

Our work operates within this framework to explore the prediction of representation in human occipitotemporal cortex, using a diverse array of open-source DNN models and a large-scale sampling of brain responses to natural images, related using model-to-brain linking methods that have become the de facto standard in the field. Surprisingly, many of the controlled comparisons among qualitatively different models yield only very small differences in their predictivity of high-level visual cortical responses. For example, networks with qualitatively different computational architectures (e.g. convolutional neural networks and vision transformers) yield almost indistinguishable brain-predictivity scores. Furthermore, networks trained to represent natural images using vision-language constrastive learning versus purely visual contrastive learning show remarkably similar capacity to predict OTC responses.

Instead, our analyses point to the importance of a model’s visual experience (i.e. input diet) as a key determinant of downstream brain predictivity that is particularly evident in the poor performance of the indoor-only-trained taskonomy models, and the high performance of OpenAI models trained on their 400M proprietary image-text database. These results indirectly reveal a currently unquantified factor of *dataset diversity* as an important predictor of brain-like visual representation. In addition, our work highlights a critical need to re-examine our standard linking assumptions and model-to-brain pipelines, potentially reducing their flexibility in order to better draw out the representational differences. We next discuss each of these results in turn, and highlight the limitations of the current approach alongside directions for future work in neuroconnectionist-style model-to-brain mapping.

### The Importance of Visual Experience

A number of our analyses point to the impact of visual input – not in the size of the image database, but in the diversity of image content – on a model’s emergent brain predictivity. First, we found that the single biggest effect was that of training: untrained models with no visual experience were unable to capture the rich representational structure evident in the late stages of the visual system. Second, impoverished diets (e.g. only faces) yield substantially lower capacity to predict brain responses than richer diets. Taskonomy models showed uniformly lower brain predictivity across the board, which we attribute to an image diet consisting of only indoor scenes. Indeed, many of the tasks in Taskonomy that seem to predict visual cortex rather poorly (e.g. semantic segmentation) seem perfectly capable of producing brain-predictive representations when trained on more diverse image sets (e.g. as is the case with the Detectron models, which rank amongst the most predictive models in our survey more generally). While there has been significant interest in the Taskonomy models for controlled model comparisons (including by us; Conwell et al., 2021; Wang et al., 2019; Dwivedi et al., 2021; Cadena et al., 2022), a direct implication of these results is that these models should not be used in future research to make arguments about which brain regions are best fit to which tasks.

The effects of increasing dataset diversity or richness beyond ImageNet1k on brain predictivity are relatively small and difficult to isolate given currently available models and datasets, but perhaps still evident. For example, we did not find clear improvements between Imagenet1K and 21K, but did find that OpenAI’s 400 million image set seems to be an important factor in positioning the CLIP models as the most predictive of all the models we surveyed (see also Wortsman et al., 2022). As a field, computational cognitive neuroscience currently has no single satisfying measure of dataset diversity or richness. Structured image similarity metrics (e.g. SSIM; Wang et al., 2004; Dosselmann and Yang, 2011) and perceptual losses (such as those behind neural style-transfer; Gatys et al., 2017; Jing et al., 2019) are in some sense early attempts to address the problem of capturing image similarity in latent representational spaces more complex than pixel statistics, but these algorithms have not been applied directly to the problem of characterizing the intrinsic richness of a visual diet. Key to solving this issue may actually be recent attempts in the computer vision community to distill smaller, less redundant, and more efficient training data from larger image sets by way of “semantic deduplication” – the removal of images from large corpora that image-aligned models like CLIP embed as effectively the same point in their latent spaces (Abbas et al., 2023) – or, similarly, through targeted pruning (Sorscher et al., 2022b). The success of semantic deduplication suggests cognitive neuroscience may be able to leverage comparable measures for understanding the minimal set of visual inputs or experiences necessary for recapitulating the overall patterns of brain responses to natural images.

Finally, another hidden factor of variation related to the input diet is not just the images themselves, but also the suite of data augmentations and related hyperparameters applied over these images during training (Wightman et al., 2021). For example, some models experience the Imagenet1K input with a ’standard’ set of augmentations which crop and rescale an image each time, with flipped left-right variation and color variation. Other models experience these same images, but with additional augmentations, leveraging some of the newer techniques that blend images and interpolate their labels (Zhang et al., 2017; Yun et al., 2019) increasing performance over standard augmentation schemes (Liu et al., 2022; Wightman et al., 2021). Relatedly, some training regimes use progressive resizing, beginning with small blurred images that ramp in resolution over training (Team, 2021; Leclerc et al., 2023). These hidden hyperparameters influence what kind of image data is fed to the model, and when, and may by implication also be important for emergent brain-predictive representation. More generally, there is a need to further curate (or build) sets of DNNs with controlled variation in image diet, image-augmentation scheme, and training recipe to explore these factors further.

### Model-to-Brain Linking Methods

In the present work we considered two different model-to-brain linking pipelines. Considered purely in isolation, our cRSA metric might lead us to conclude that our most predictive model layers explain less than a third (*∼*31%) of the explainable variance in the Natural Scenes Dataset; conversely, our veRSA metric would suggest we have explained the vast majority of that variance (*∼*79.5%). Currently, it is somewhat unclear which of these is the more correct, not least because we have yet to cohere as a field on the principles of mechanistic intepretability in modeling that logically favor one over the other (Kay, 2018; Cao and Yamins, 2021; Han et al., 2023). Relatedly, our model-to-model comparisons highlight that models trained with different inductive biases are indeed learning different representational formats, as revealed by classical RSA. However, the feature re-weighting procedure effectively solves for a similar subspace present in all of the learned feature spaces. This difference demonstrates that there is meaningful diversity in the learned representations of models that our metrics are failing to translate into significantly different brain-predictivity scores. These observations lead to several possible directions for stronger, more diagnostic model-to-brain comparisons.

One possible way forward is to develop deeper theoretical commitments to the relationship between single units in the model and single neurons or voxels. For example, adding a sparse positive regularization term to our linear encoding models might better capture the functional role of model unit tuning and require more aligned tuning curves (Prince and Konkle, 2023). Relatedly, we could consider different commitments on the coverage between a given set of model units and brain targets: Perhaps we should allow for the selection of units from across multiple model layers to better account for differences in representational hierarchy (Nonaka et al., 2021), or maybe even explore one-to-one mappings that require single units or features in models to directly predict single units in the brain (Arend et al., 2018)? Finally, we could expand the brain target to include not just predictivity of the regional geometry and single unit or voxel tuning, but also its topographic organization (Lee et al., 2020; Blauch et al., 2022; Margalit et al., 2023; Doshi and Konkle, 2023).

Another possible route forward is to more carefully select the set of images from which both brain data and model responses are collected. The Natural Scenes Dataset (Allen et al., 2022) contains reliably measured brain responses to a wide variety of natural images – providing clear generality over prior fMRI datasets that have focused on isolated objects over a white background. However, our results highlight that, with this widely sampled natural image set, basically all performant models can capture the major large-scale image distinctions present in the visual system responses. One interpretation of this finding is that this widely sampled image set may actually be obscuring finer-scale representational differences that could be captured by some, but not all, of these models. For example, one relevant approach formulated to address this limitation has been that of ‘controversial stimuli’ (Golan et al., 2020, 2022), a generative method that leverages gradient ascent techniques to create novel synthetic images that actively differentiate one model from another. A related idea that could be applied to existing datasets is ‘controversial selection’ – a model-informed, data-driven selection of natural images that maximizes the difference in direct model-to-model comparisons independently of model-to-brain comparisons, and in a way that is more agnostic to the models themselves. Finally, it is possible that a return to more artificial stimuli (e.g. line drawings, or geometric shapes) or a small number of key diagnostic psychophysical comparisons (e.g. upright versus inverted, or scrambled faces) could provide complementary fine-grained insight to further differentiate models in their ability to predict not just visual brains, but visual behavior (Rust and Movshon, 2005; Krakauer et al., 2017; Bowers et al., 2022).

### Relationship to Prior Work

Our approach has many similarities to neural benchmarking endeavors, aimed at identifying the most brain-like model of the visual system, whether in primates or mice (Schrimpf et al., 2018a, 2020; Cichy et al., 2019; Willeke et al., 2022). This approach typically focuses on an aggregate ‘brain-score’, which involves scoring models on data from multiple hierarchical regions, across multiple datasets, and ranking models on a leaderboard according to their aggregate scores. Here, we took a complementary, but notably different approach, leveraging the variation in a large sample of diverse open-source models to draw insights not from which individual models are best, but from the underlying factors that define the average performance of a distinct *set* of models. Moreover, our analysis focused solely on the structure of the population code of the late-stages of the ventral visual stream (OTC), motivated by the fact that the format of this code has been a somewhat elusive target in neuroscience, with extensive speculation about the pressures shaping its formation (Hasson et al., 2002; DiCarlo and Cox, 2007; Op de Beeck et al., 2008; Mahon and Caramazza, 2011; Arcaro and Livingstone, 2021; Konkle and Alvarez, 2022). Thus, our aim was to focus on how major variations in architecture, task, and visual diet – three crucial elements of the neuroconnectionist research programme – give rise to different learned features and resulting late-stage ventral visual system predictivity.

Our results contrast with some of the emerging concurrent work in this domain, e.g. arguing for effective dimensionality (Elmoznino and Bonner, 2022), or the primacy of language-aligned features (Wang et al., 2022) as sources of increased brain predictivity. For example, our work does not provide immediate support for effective dimensionality as a model-agnostic predictor of brain-like representation. Importantly, however, we do not consider this a claim on the importance of similar model-agnostic metrics more generally: the intersection of manifold statistics and neural coding schemes is an emerging field (Cohen et al., 2020; Sorscher et al., 2022a; Yerxa et al., 2023), which we believe will be fruitful for the neuroconnectionist research program, providing deeper insights into representation learning in brains and models alike. Relatedly, another divergent result from our work suggests that language alignment is not in fact the key pressure governing the superlative performance of the OpenAI-CLIP models. Here, too, however, we note that there remains substantial room for more targeted analyses at finer-grained neural resolution and with targeted stimuli that emphasize vision-only versus language-aligned models. These kinds of analyses will be crucial for elucidating what are hypothesized to be significant interactions between vision and language at the most anterior portions of the ventral stream (Popham et al., 2021; Tang et al., 2023). Indeed, one limitation of our analysis in its current form is that differences in predictivity in smaller (sub)regions of occipitotemporal cortex may potentially be obscured in our more general mask.

More broadly, even with the 1.8 billion regressions and 50.3 thousand representational similarity analyses undergirding the results reported in this paper, we note that our survey approach reflects only a small slice of the possibility space for model-to-brain comparisons with this dataset. For example, we did not try to link hierarchical model layers with hierarchical brain regions (e.g. Nonaka et al., 2021), or focus in on category-selective regions (e.g. Ratan Murty et al., 2021; Prince and Konkle, 2020) and early visual cortex (though see Supplementary Information SI.4 for initial analyses). To facilitate future work on these fronts, we have open-sourced our highly optimized DeepDive code package (see Code Availability Section), which can be used to conduct these model-to-model, and model-to-brain analyses at scale.

Our aim here was to provide a broad ‘lay-of-the-land’ for relating deep neural network models to the high-level visual system – leveraging controlled variation present in the diversity of available models to conduct opportunistic experiments that isolate factors of architecture, task, and dataset (Muttenthaler et al., 2022; Doerig et al., 2023; Kanwisher et al., 2023; Ren and Bashivan, 2023). Taken together, our results call for a deeper investigation into the impact of visual diet diversity, and highlight the need for conceptual advances in developing theoretically-constrained linking procedures that relate models to brains, along with more diagnostic image sets to further differentiate these highly performant computer vision models.

## Methods

### Model Selection

We collected a set of 224 distinct models (160 trained; 64 randomly-initialized), sourced from the following repositories: the Torchvision (PyTorch) model zoo (Paszke et al., 2019); the Pytorch-Image-Models (timm) library (Wightman, 2019); the VISSL (self-supervised) model zoo (Goyal et al., 2021); the OpenAI CLIP collection (Radford et al., 2021); the PyTorch Taskonomy (visualpriors) project (Zamir et al., 2018; Sax et al., 2018, 2019); the Detectron2 model zoo (Wu et al., 2019); and Harvard Vision Sciences Laboratory’s Open-IPCL project (Konkle and Alvarez, 2022).

This set of models was collected with a focus on high-level visual representation, and was explicitly intended to span different architectural types, training objectives, and other available variations. Our 64 randomly-initialized models consist of the untrained variants of each ImageNet-1K-trained architecture from the Torchvision and Pytorch-Image-Models repository. (These models were initialized using the default parameters provided by the package, and in most cases are the recommended defaults specified by the contributing authors). Where possible, we extracted the relevant metadata for each model using automatic parsing of web data from the associated repositories; where this automatic parsing was not possible, we manually annotated each model with respect to its associated publication. A list of all included models and their most significant metadata (architecture, task, training data) is included in SI Table 1.

### Human fMRI Data

The Natural Scenes Dataset (Allen et al., 2022) contains measurements of over 70,000 unique stimuli from the Microsoft Common Objects in Context (COCO) dataset (Lin et al., 2014) at high resolution (7T field strength, 1.6-s TR, 1.8*mm*^3^ voxel size). In this analysis, we focus on the brain responses to 1000 COCO stimuli that overlapped between subjects, and limit analyses to the 4 subjects (subjects 01, 02, 05, 07) that saw these images in each of their 3 repetitions across scans. The 3 image repetitions were averaged to yield the final voxel-level response values in response to each stimulus. All responses were estimated using a new, publicly available GLM toolbox (GLMsingle; Prince et al., 2022), which implements optimized denoising and regularization procedures to accurately measure changes in brain activity evoked by experimental stimuli.

### Voxel Selection Procedure

To achievable a reasonable signal-to-noise ratio (SNR) in our target data, we implement a reliability-based voxel selection procedure (Tarhan and Konkle, 2020) to subselect voxels containing stable structure in their responses. Specifically, we use the NCSNR (“noise ceiling signal-to-noise ratio”) metric computed for each voxel as part of the NSD metadata (Allen et al., 2022) as our reliability metric. In this analysis, we include only those voxels with NCSNR > 0.2.

After filtering voxels based on their NCSNR, we then filtered voxels based on region-of-interest (ROI). In our main analyses, we focus on voxels within occipitotemporal cortex (OTC; also referred to as human IT). Our goal was to identify a sector of cortex beyond early visual cortex that covers the ventral and lateral object-responsive cortex, including category-selective regions. To do so, we first considered voxels within a liberal mask of the visual system (“nsdgeneral” ROI). Next we selected the subset within either the mid-to-high ventral or mid-to-high lateral ROIs (“streams” ROIs). Then, we included all voxels from 11 category-selective ROIs (face, body, word, and scene ROIs, excluding RSC) with a *t*-contrast statistic > 1; while many of these voxels were already contained in the streams ROIs, this ensures that these regions were included in the larger scale OTC sector. The number of OTC voxels included were 8,088 for subject 01, 7,528 for subject 02, 8,015 for subject 05, and 5,849 for subject 07, for a combined total of 29,480 voxels.

### Noise Ceilings

To contextualize model performance results, we estimated noise ceilings for each of the target brain ROIs. These noise ceilings indicate the maximum possible performance that can be achieved given the level of measurement noise in the data. Importantly, in the present context, noise ceiling estimates refer to within-subject representational similarity matrices (RSMs), where noise reflects trial-to-trial variability in a given subject. This stands in contrast to more conventional group-level representational dissimilarity matrices (Kriegeskorte et al., 2008a), where noise reflects variability across subjects. To estimate within-subject noise ceilings, we developed a novel method based on generative modeling of signal and noise, which we term GSN (https://github.com/cvnlab/GSN/). This method estimates, for a given ROI, multivariate Gaussian distributions characterizing the signal and the noise under the assumption that observed responses can be characterized as sums of samples from the signal and noise distributions. A post-hoc scaling is then applied to the signal distribution such that the signal and noise distributions generate accurate matches to the empirically observed reliability of RSMs across independent splits of the experimental data. Noise ceilings are estimated using Monte Carlo simulations in which a noiseless RSM (generated from the estimated signal distribution) is correlated with RSMs constructed from noisy measurements (generated from the estimated signal and noise distributions). All noise ceiling calculations were performed on independent data outside the main analysis.

### Feature Mapping Methods

#### Feature Extraction Procedure

For each of our candidate DNN models, we extract features in response to each of our probe stimuli at each distinct layer of the network. Importantly, we define a layer here as a distinct computational (sub)module. This means, for example, that we treat convolution and the rectified nonlinearity that follows it as two distinct feature maps; crucially, for transformers, this also means we analyze the outputs not only of each attention head, but of the individual key-query-value modules used to compute them. At the end of our feature extraction procedure, for each model and each model layer, we arrive at a feature matrix of dimensionality number-of-images x number-of-features, the latter value of which represents the flattened dimensions of the original feature map. Beyond flattening, we perform no other transformation of the original features during extraction.

#### Classical RSA (cRSA)

To compute the classical representational similarity (cRSA) score (Kriegeskorte et al., 2008a) for a single layer, we used the following procedure: First, we split the 1000 images into two sets of 500 (a training set, and a testing set). Using the training set of images, we compute the representational similarity matrices (RSMs) of each model layer (500 x 500 x number-of-layers) using Pearson correlation as the distance metric. We then compare each layer’s RSM to the brain RSM, also using Pearson similarity, and identify the layer with the highest correlation as the model’s most brain-predictive layer. Finally, using the held-out test set of 500 images, we compute that target layer’s RSM and correlate it with the brain RSM. This test score from the most predictive layer serves as the overall cRSA score for the target model.

#### Voxel-Encoding RSA (veRSA)

To arrive at a voxel-encoding representational similarity (veRSA) score (Khaligh-Razavi et al., 2017b; Kaniuth and Hebart, 2022; Konkle and Alvarez, 2022) for a single model, the overall procedure was similar to that of cRSA, but with the addition of an intermediate encoding procedure wherein layerwise model features were fit to each individual voxel’s response profile across the image probe set.

The first step in the encoding procedure is the dimensionality reduction of model feature maps. We perform this step for two reasons: first, the features extracted from various deep neural networks can sometimes be massive (the first convolutional layer of VGG16, for example, yields a flattened feature matrix with *∼*3.2 million dimensions per image); and second, the same dimensionality reduction procedure applied to all layers ensures that the explicit degrees of freedom across model layers is constant. To reduce dimensionality, we apply the SciKit-Learn implementation of sparse random projection (Pedregosa et al., 2011). This procedure relies on the Johnson-Lindenstrauss (JL) lemma (Achlioptas, 2001), which takes in a target number of samples and an epsilon distortion parameter, and returns the number of random projections necessary to preserve the euclidean distance between any two points up to a factor of 1*±*epsilion. (Note that this is a general formula; no brain data enter into this calculation). In our case, with the number of samples set to 1000 (the total number of images) and an epsilon distortion of 0.1, the JL lemma yields a target dimensionality of 5920 projections.

After computing this target dimensionality, we then proceed to compute the sparse random projection for each layer of our target DNN. The sparse random projection matrix consists of zeros and sparse ones of nearly orthogonal dimensions, normalized by the density of the matrix (the inverse square root of the total number of features). The layerwise feature maps are then projected onto this matrix by taking the dot product between them. The output of the procedure is a reduced layerwise feature space of size of 1000 images x 5920 dimensions with a preserved representational geometry. Note that in cases where the number of features is less than the number of projections suggested by the JL lemma, the original feature map is effectively upsampled through the random projection matrix, again yielding a matrix of 1000 x 5920 dimensions.

We compute our encoding model for each voxel as a weighted combination of these 5920 dimensions, using brain data from our training set of 500 images. (We note that while the number of dimensions needed for only 500 images would be only D = 5326 according to the JL lemma, adding extra dimensions will only preserve the geometry with nominally less distortion than the epsilon provided, and does not meaningfully affect the results). The fitting procedure for each voxel leverages SciKit-Learn’s cross-validated ridge regression function (“RidgeCV”), a hyperefficient regression method that uses generalized cross-validation to provide a LOOCV prediction per image (per output). This fit was computed over a logarithmic range of alpha penalty parameters (1*e^−^*^1^ to 1*e^−^*^7^), to identify each voxel’s optimal alpha parameter. We modified the RidgeCV function in order to select the best alpha using Pearson correlation as a score function (the same score function we use to evaluate the model at large), and to parallelize an internal for-loop for greater efficiency. This yielded a set of encoding weights for each voxel (number-of-voxels x 5920 reduced-feature-dimensions).

Next, with these encoding weights and the 500 training images, we compute the predicted response of every voxel to each image, and compute the corresponding *predicted* RSM using Pearson correlation. After computing each layer’s representational similarity via Pearson correlation between the layer-predicted RSM and the target brain RSM, we again select the most predictive layer on the basis of results from the training set and compute this layer’s RSA correspondence to the brain data using the held-out set of 500 test images. This test score from the most predictive layer serves as the final veRSA score for the target model.

We emphasize that this method contrasts with popular practices in primate and mouse benchmarking, which treat predictivity of unit-level univariate response profiles as the key outcome measure. Because fMRI affords more systematic spatial sampling over the cortex, rather than taking the aggregate of single voxel fits as our key measure, we choose to treat the population representational geometry over each ROI as the critical target for prediction. This multi-voxel similarity structure provides different kinds of information about the format of population-level coding than do individual units (Kriegeskorte et al., 2008b). Computing the veRSA metric does, however, yield individual voxelwise-encoding models, the individual predictive accuracies of which we collect and have available in addition to the cRSA and veRSA scores for future analysis.

### Statistical Analysis

#### Opportunistic Experiments

The statistical test for each targeted model set comparison consisted of a linear fixed effects model with brain-predictivity score (cRSA or veRSA, in units of *r_P earson_*) as the outcome variable, and two additive effects: that of the experimental manipulation and that of subject ID (to control for overall differences in predictivity across subjects). In the case of the Imagnet1K versus Imagenet21K comparison, a mixed effects model was used to capture the within-model comparison structure of the comparison, with an additional random intercept for model ID. All linear fixed and mixed effects models were fit using R’s ‘stats’ (R Core Team, 2013) and ‘lme4’ (Bates et al., 2015) packages, respectively. Confidence intervals and p-values were estimated using R’s ‘parameters’ package (Lüdecke et al., 2020), which leverages a Wald *t*-distribution approximation (for fixed effects), and a Wald *z*-distribution approximation (for random effects).

We provide each of these linear models (in R-style formulaic pseudocode) below:

CNNs versus Transformers:

lm(*Score ∼ ArchitectureClass* + *SubjectID,* reference = CNN)

Taskonomy Encoders:

lm(*Score ∼ Task* + *SubjectID,* reference = Denoising)

Contrastive Self-Supervised Learning:

lm(*Score ∼ TaskClass* + *SubjectID,* reference = Non-Contrastive)

Language Alignment:

lm(*Score ∼ TaskClass* + *SubjectID,* reference = SimCLR)

ImageNet1K versus ImageNet21K:

lmer(*Score ∼ DatasetSize* + *SubjectID* + (1*|ModelID*), reference = ImageNet1K)

Objects, Faces, Places:

lm(*Score ∼ Dataset* + *SubjectID,* reference = ImageNet1K-Objects)

#### Breakpoint Analysis of Model Rankings

When analyzing variation across all models, we use a breakpoint analysis (Muggeo, 2003) to quantify the distinct visual elbows that appear in the rank-order of these models. This piece-wise regression method estimates the origin and terminus of distinct linear sub-segments across an otherwise nonlinear trend. Here, we perform this analysis using R’s ‘segmented’ package (Muggeo, 2008), predicting score by model rank, setting the sole *ψ* hyperparameter (N*_Breakpoints_*) to 2, and computing confidence intervals with the ‘gradient’ method (Muggeo, 2017) for breakpoint interval estimation. We use the first breakpoint yielded by this analysis (over the veRSA brain-predictivity scores) to divide the full set of models into the higher-ranking (‘top’) and lower-ranking (‘bottom’) model sets that we use in several subsequent analyses.

#### Model-to-Model Analysis

To compare the representations of our surveyed models directly, we perform a model-to-model representational similarity analysis, comparing the RSMs from the most brain-predictive layer of each surveyed model directly, with (as in cRSA) or without (as in veRSA) the intermediate encoding of voxels. As in our main analyses, we compute these RSMs with a first-order Pearson correlation, then compare their flattened upper triangular portions with a second-order Pearson correlation.

We reduce the dimensionality (for visualization’s sake) of the resultant modelwise-RSM to 2 dimensions with classical multidimensional scaling (MDS; Gower, 1966), as implemented in R’s ‘stats’ package (R Core Team, 2013). To better visualize the representational similarities of distinct groups of models in this reduced space, we draw convex hulls around each group using R’s ‘ggforce’ package (Pedersen, 2022), with the relative concavity of each hull set to 10.

#### Effective Dimensionality

We compute the effective dimensionality (ED) of each of our target layers using the formula proposed by Del Giudice (2021): the squared sum of all eigenvalues, divided by the sum of the squared eigenvalues.

Note that we compute the ED of these representations both with and without sparse random projection. In a noteworthy validation of the JL lemma’s preservation of pointwise geometry, we find no analytically-relevant difference in the measured value of ED as a function of this dimensionality reduction (*r_Spearman_* = 0.993 between ED with and without SRP, across targeted layers). Given this minimal difference, we use the effective dimensionality of the randomly-projected representations in all main analyses, as these were the feature spaces that were directly used in predicting the brain.

Our approach does differ from that of Elmoznino and Bonner (2022), as we perform no pooling or other forms of feature aggregation on our targeted layers before computing the ED of these layers. See Supplementary Information SI.2 for more detailed comparisons. We choose not to do a pooling operation before calculating effective dimensionality because (1) this is not a general operation that can be performed on other kinds of (non-convolutional) architectures, and (2) estimating the effective dimensionality over the same feature space used to fit the brain responses seems preferable from the theoretical standpoint of establishing a measure of model variation that meaningfully abstracts over the details of model implementation.

## Acknowledgments

This work was supported by a Hodgson’s Innovation Fund Grant (C.C.), an NDSEG fellowship (J.S.P.), and NSF-CAREER BCS-1942438 (T.K). Collection of the NSD dataset was supported by NSF IIS-1822683 (K.K.) and NSF IIS-1822929.

## Supplementary Information

### SI.1 Surveyed Models

The list of all surveyed models may be found in Table 1 below.

### SI.2 Effective Dimensionality

Concurrent work by Elmoznino and Bonner (2022) found that model feature spaces with greater effective dimensionality (ED) were better predictors of high-level visual cortex, including in prediction of occipitotem-poral responses in the Natural Scenes Dataset. In our analysis, we do not find this trend. What reasons might account for this disjunct between findings?

First, we considered model sets. At the time of this preprint, the models tested by Elmoznino and Bonner (2022) include only ResNet18 and ResNet50 models, trained via self-or category-supervision on ImageNet1K; the ResNet50 models of Taskonomy; and untrained ResNet18 and ResNet50 models. When we subsetted our models to include only the best layers of the Taskonomy Resnet50s (N = 24), we found that ED was indeed a significant positive predictor of OTC predictivity (*r_Spearman_* = 0.41 [0.015, 0.73], *p* = 0.026 in cRSA; 0.435 [0.162, 0.730], *p* = 0.039 in veRSA). However, as is perhaps already evident from the large confidence intervals, we found this effect to be somewhat brittle, quickly breaking with the addition of the 12 less-performant models that ranked similarly to those of Taskonomy (and below the breakpoint suggested by the segmented regression analysis; *r_Spearman_* = 0.041 [-0.357, 0.392], *p* = 0.341 in cRSA; 0.0532 [-0.324, 0.393], *p* = 0.37 in veRSA).

Second, we tested the impact of the image set on computation of ED. Elmoznino and Bonner (2022) estimated ED over 10,000 ImageNet1K validation images, whereas we estimated ED over the ‘shared1000’ COCO images. To check for differences in ED estimates computed over the different image sets, we directly replicated the approach of Elmoznino and Bonner (2022). This involved considering only the ReLU stages of each convolutional block, and performing a global-pooling operation over features of each model layer, prior to computing ED. While we did not include this pooling step in our main analyses, we did so in this supplementary analysis to ensure that ED levels across the two image sets could be compared with all other analytical choices held constant. The results of this analysis can be seen in Supplementary Figure SI.1. We observe similar ED estimates when probing these two different image sets, with slightly lower ED estimates in the later layers when using the ‘shared1000’ probe set.

**Figure SI.1:**
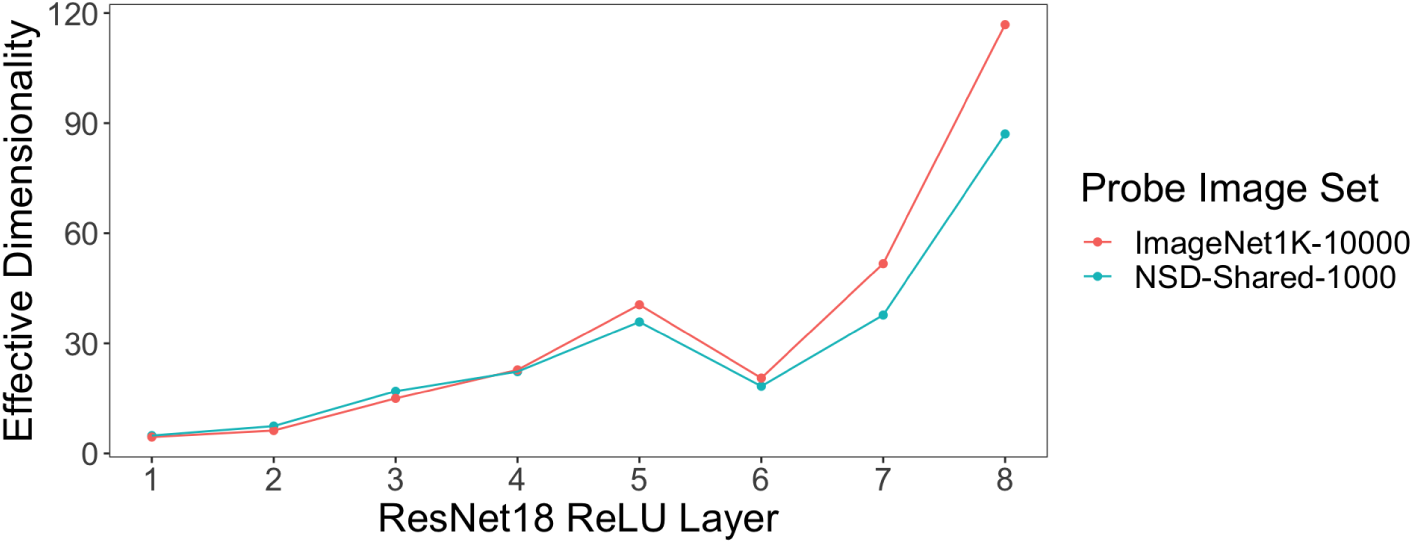
Results from a direct replication of Elmoznino and Bonner (2022)’s quantification of effective dimensionality (ED). We use the ReLU outputs of ResNet18’s residual blocks to determine whether the ‘shared1000’ COCO images were a sufficiently diverse image set for calculating a meaningful measure of ED, compared with 10,000 ImageNet1K validation used in previous analyses. While the estimated ED yielded by the 10,000 ImageNet1K images is visibly higher in later layers, the correlation between the ED across layers (which undergirds the primary statistics we use to assess the relationship of ED to brain predictivity in our main analysis) is effectively 1:1 (*r_P earson_* = 0.99 [0.98, 0.99]). This shows that the ‘shared1000’ COCO images are a sufficiently diverse image set for calculating ED.

Broadly, there are several key analytical differences in our test of ED and brain predictivity that likely underlie the divergence between our findings and those of Elmoznino and Bonner (2022). First, Elmoznino and Bonner (2022) jointly analyzed ED from multiple layers of individual models, as well as from untrained models. We argue that this choice introduces significant covariation between high-level feature quality and effective dimensionality. If the importance of ED for high brain predictivity was truly a general principle, then our stronger test examining the ED variation across models from only the most brain-aligned layer should also yield correlations with brain predictivity (but this was not the case). Second, Elmoznino and Bonner (2022) applied a global max-pooling to the convolutional feature maps before computing ED, noting their primary interest in the variance of image features, rather than the variance in those properties across space. However, in our analysis pipeline, such a global max-pooling operation is frequently infeasible (e.g, for non-convolutional models). And, more generally, estimating ED over the exact same feature space used to fit the brain responses seems preferable from a theoretical standpoint.

### SI.3 Stability of Rankings across Subjects

In our main analysis, all predictivity scores we report are the average across 4 subjects. That even the smallest differences in predictivity between models is significant across many of our controlled model comparisons already suggests that patterns in these scores are meaningfully stable and consistent across subjects. We further quantity this stability in two ways.

The first is with a rank-order correlation of model rankings between individual pairs of subjects (N = 6 unique pairwise comparisons). Across *all models* – including randomly-initialized models – this correlation is *r_Spearman_* = 0.98 [0.975, 0.985] for cRSA and 0.957 [0.945, 0.967]. Across only the top 125-ranking models, this correlation remains remarkably high, at *r_Spearman_*= 0.923 [0.907, 0.938] for cRSA and *r_Spearman_* = 0.835 [0.797, 0.873] for eRSA. A permutation test in which we scramble the model scores across subjects and recompute this same rank order correlation suggests these values are *extremely* unlikely to occur by chance (mean permuted *r_Spearman_*= 0.0052 [-0.0455, 0.0736] in cRSA and 0.008 [-0.0239, 0.0278] in eRSA; not a single permutation achieves *r_Spearman_*> 0.1).

What we are testing here, in effect, is the logic of brain-predictivity leaderboards. This analysis suggests that – while small – the differences between consecutively ranked models in our analyses are statistically meaningful. This does not spare inferences derived from leaderboards from the critique that differences between consecutively ranked models are rarely attributable to unique or controlled sources of variation, but it does suggest that small score differences reflect more than random statistical variation.

### SI.4 Modeling Results in Early Visual Cortex

While the focus of our main analyses was the predictivity of a unified OTC ROI, we designed our pipeline to generate predictions for a number of additional ROIs – including early visual cortex (EVC: V1-V4).

This ROI encapsulates the ventral and dorsal aspects of areas V1, V2, and V3, as well as area hV4. To define the EVC ROI for each subject, we again first isolated voxels within the “nsdgeneral” ROI, and then selected for analyses any voxels that both fell within one of the early visual regions listed above, and that exceeded the NCSNR threshold of 0.2. This procedure yielded a total of 4,657 voxels for subject 01, 3,757 voxels for subject 02, 3,661 voxels for subject 05, and 3,251 voxels for subject 07.

A first question we might ask, then, is whether the better models of OTC in general are better models of EVC. To do this, we can correlate the predictivity of each model we test in OTC with that same model’s predictivity in a macro-scale EVC ROI. (Note that, as in our analyses of OTC, we select the most EVC-predictive layer from each model using the training set of 500 images, and report the score of this layer on the 500 held-out test images). Across *all models*, this correlation is high: *r_Spearman_*= 0.809 [0.801, 0.822] in cRSA and 0.835 [0.816, 0.853] in eRSA. Across only the 125 highest ranking models of OTC, this correlation is markedly lower: 0.212 [0.166, 0.261] in cRSA and 0.287 [0.209, 0.364] in eRSA. In other words, it seems, poor models of OTC (e.g. the Taskonomy-trained and randomly-initialized) models are also poor models of EVC; excluding these poor-performing models, however, better models of OTC are not necessarily better models of EVC.

This rank-order correlation across many models, more generally, does not necessarily capture the subtleties and trends we saw in our opportunistic experiments and controlled model comparisons, which we can directly repeat in EVC. While a comprehensive recap of each opportunistic experiment applied to EVC is beyond the scope of this analysis, what we can say is that many, but not all, of the trends we observe in OTC are recapitulated in EVC. For example, there is, once again, only a negligible difference in the average predictivity of CNNs versus transformers (cRSA *β* = -0.029, p = 7.08e-10; veRSA *β* = -0.007, p = 2.61e-3). Perhaps the most notable divergence between EVC and OTC in terms of our opportunistic experiments is the minimal difference of the veRSA metric in EVC between the self-supervised (IPCL) model trained on objects (ImageNet) versus faces (VGGFace2) (EVC *β* = -0.0394, p = 8.62e-4, OTC *β* = -0.2719, p = 1.03e-9). Unlike in OTC, then, face-trained models (with reweighting) perform on par with object-trained models. This means that even the depleted set of natural image statistics available in a visual diet of faces alone may still be sufficient to capture core aspects of early visual cortical representation.

### SI.5 Compute Required

We used a single machine with 8 Nvidia RTX 3090 GPUs, 755gb of RAM, and 96 CPUs. GPUs were used only for extracting model activations, and could (without major slowdown) be removed from the analytic pipeline. Dimensionality reduction and regression computations were CPU and RAM intensive. Replicating all of our results would take approximately three weeks on a similar machine.

## Footnotes

2 Analysis by the Taskonomy authors suggests that only 100 of the 1000 imagenet classes are present across the ’scenes’ of the Taskonomy dataset (Zamir et al., 2018) – suggesting this difference may be partially attributable to impoverished category variation.

3 Note that a more precise characterization of this trend actually indicates *two* visual elbows: the first (as outlined above) putatively corresponds to *training diet*, and the second putatively corresponds to the role of *training itself*, since it roughly demarcates the upper bound of brain predictivity for the randomly-initialized models. We thus parametrized our segmented regression analysis to fit for two breakpoints (*ψ*). These 2 *ψ* were equal to 124 [123.6, 124.7] and 154 [153.7, 154.9] in veRSA, and ranks 123 [122.31, 124.34] and 150 [149.2, 151.3] in cRSA. (For further details, see Methods).

## Notes

### Competing Interest Statement

The authors have declared no competing interest.

### Summary of Updates

The new version of this manuscript constitutes a substantial reworking of the first, building on its core results, but adding a significant number of analyses and a new, more general framework for interpreting them.

## References

1. Abbas, A., Tirumala, K., Simig, D., Ganguli, S., and Morcos, A. S. (2023). Semdedup: Data-efficient learning at web-scale through semantic deduplication. arXiv preprint arXiv:2303.09540.

2. Achlioptas, D. (2001). Database-friendly random projections. In Proceedings of the twentieth ACM SIGMOD-SIGACT-SIGART symposium on Principles of database systems, pages 274–281.

3. Allen, E. J., St-Yves, G., Wu, Y., Breedlove, J. L., Prince, J. S., Dowdle, L. T., Nau, M., Caron, B., Pestilli, F., Charest, I., and others (2022). A massive 7T fMRI dataset to bridge cognitive neuroscience and artificial intelligence. Nature neuroscience, 25(1):116–126. Publisher: Nature Publishing Group US New York.

4. Arcaro, M. J. and Livingstone, M. S. (2021). On the relationship between maps and domains in inferotemporal cortex. Nature Reviews Neuroscience, 22(9):573–583.

5. Arend, L., Han, Y., Schrimpf, M., Bashivan, P., Kar, K., Poggio, T., DiCarlo, J. J., and Boix, X. (2018). Single units in a deep neural network functionally correspond with neurons in the brain: preliminary results. Technical report, Center for Brains, Minds and Machines (CBMM).

6. Bashivan, P., Kar, K., and DiCarlo, J. (2019). Neural population control via deep image synthesis. Science, 364.

7. Bates, D., Mächler, M., Bolker, B., and Walker, S. (2015). Fitting linear mixed-effects models using lme4. Journal of Statistical Software, 67(1):1–48.

8. Blauch, N. M., Behrmann, M., and Plaut, D. C. (2022). A connectivity-constrained computational account of topographic organization in primate high-level visual cortex. Proceedings of the National Academy of Sciences, 119(3):e2112566119.

9. Bowers, J. S., Malhotra, G., Dujmović, M., Montero, M. L., Tsvetkov, C., Biscione, V., Puebla, G., Adolfi, F., Hummel, J. E., Heaton, R. F., et al. (2022). Deep problems with neural network models of human vision. Behavioral and Brain Sciences, pages 1–74.

10. Cadena, S., Willeke, K., Restivo, K., Denfield, G., Walker, E., Sinz, F., Bethge, M., Tolias, A., and Ecker, A. (2021). A diverse task-driven characterization of early and mid-level representations of the primate ventral stream. In Computational and Systems Neuroscience Meeting (COSYNE 2021).

11. Cadena, S. A., Denfield, G. H., Walker, E. Y., Gatys, L. A., Tolias, A. S., Bethge, M., and Ecker, A. S. (2019). Deep convolutional models improve predictions of macaque V1 responses to natural images. PLoS computational biology, 15(4):e1006897. Publisher: Public Library of Science.

12. Cadena, S. A., Willeke, K. F., Restivo, K., Denfield, G., Sinz, F. H., Bethge, M., Tolias, A. S., and Ecker, A. S. (2022). Diverse task-driven modeling of macaque V4 reveals functional specialization towards semantic tasks. bioRxiv, pages 2022–05. Publisher: Cold Spring Harbor Laboratory.

13. Cadieu, C. F., Hong, H., Yamins, D. L., Pinto, N., Ardila, D., Solomon, E. A., Majaj, N. J., and DiCarlo, J. J. (2014). Deep neural networks rival the representation of primate IT cortex for core visual object recognition. PLoS computational biology, 10(12):e1003963. Publisher: Public Library of Science San Francisco, USA.

14. Cao, R. and Yamins, D. (2021). Explanatory models in neuroscience: Part 1–taking mechanistic abstraction seriously. arXiv preprint arXiv:2104.01490.

15. Chen, T., Kornblith, S., Norouzi, M., and Hinton, G. (2020). A simple framework for contrastive learning of visual representations. In International Conference on Machine Learning, pages 1597–1607. PMLR. arXiv preprint arXiv:2002.05709.

16. Cichy, R., Khosla, A., Pantazis, D., Torralba, A., and Oliva, A. (2016). Comparison of deep neural networks to spatiotemporal cortical dynamics of human visual object recognition reveals hierarchical correspondence. Scientific Reports, page 6.

17. Cichy, R. M., Roig, G., Andonian, A., Dwivedi, K., Lahner, B., Lascelles, A., Mohsenzadeh, Y., Ramakrishnan, K., and Oliva, A. (2019). The algonauts project: A platform for communication between the sciences of biological and artificial intelligence. arXiv preprint arXiv:1905.05675.

18. Cohen, U., Chung, S., Lee, D. D., and Sompolinsky, H. (2020). Separability and geometry of object manifolds in deep neural networks. Nature communications, 11(1):746.

19. Conwell, C., Mayo, D., Barbu, A., Buice, M., Alvarez, G., and Katz, B. (2021). Neural regression, representational similarity, model zoology & neural taskonomy at scale in rodent visual cortex. Advances in Neural Information Processing Systems, 34:5590–5607.

20. Dehghani, M., Djolonga, J., Mustafa, B., Padlewski, P., Heek, J., Gilmer, J., Steiner, A., Caron, M., Geirhos, R., Alabdulmohsin, I., and others (2023). Scaling vision transformers to 22 billion parameters. arXiv preprint arXiv:2302.05442.

21. Del Giudice, M. (2021). Effective dimensionality: A tutorial. Multivariate behavioral research, 56(3):527–542. Publisher: Taylor & Francis.

22. DiCarlo, J., Zoccolan, D., and Rust, N. (2012). How Does the Brain Solve Visual Object Recognition? Neuron, 73(3):415–434.

23. DiCarlo, J. J. and Cox, D. D. (2007). Untangling invariant object recognition. Trends in cognitive sciences, 11(8):333–341.

24. Doerig, A., Sommers, R. P., Seeliger, K., Richards, B., Ismael, J., Lindsay, G. W., Kording, K. P., Konkle, T., Van Gerven, M. A., Kriegeskorte, N., et al. (2023). The neuroconnectionist research programme. Nature Reviews Neuroscience, pages 1–20.

25. Doshi, F. R. and Konkle, T. (2023). Cortical topographic motifs emerge in a self-organized map of object space. Science Advances, 9(25):eade8187.

26. Dosselmann, R. and Yang, X. D. (2011). A comprehensive assessment of the structural similarity index. Signal, Image and Video Processing, 5:81–91.

27. Dwivedi, K., Bonner, M. F., Cichy, R. M., and Roig, G. (2021). Unveiling functions of the visual cortex using task-specific deep neural networks. PLoS computational biology, 17(8):e1009267. Publisher: Public Library of Science San Francisco, CA USA.

28. Eickenberg, M., Gramfort, A., Varoquaux, G., and Thirion, B. (2017). Seeing it all: Convolutional network layers map the function of the human visual system. NeuroImage, 152:184–194. Publisher: Elsevier.

29. Elmoznino, E. and Bonner, M. F. (2022). High-performing neural network models of visual cortex benefit from high latent dimensionality. bioRxiv, pages 2022–07. Publisher: Cold Spring Harbor Laboratory.

30. Gallicchio, C. and Scardapane, S. (2020). Deep randomized neural networks. In Recent Trends in Learning From Data: Tutorials from the INNS Big Data and Deep Learning Conference (INNSBDDL2019), pages 43–68. Springer.

31. Garrido, Q., Balestriero, R., Najman, L., and Lecun, Y. (2022). Rankme: Assessing the downstream performance of pretrained self-supervised representations by their rank. arXiv preprint arXiv:2210.02885.

32. Gatys, L. A., Ecker, A. S., Bethge, M., Hertzmann, A., and Shechtman, E. (2017). Controlling perceptual factors in neural style transfer. In Proceedings of the IEEE conference on computer vision and pattern recognition, pages 3985–3993.

33. Geirhos, R., Narayanappa, K., Mitzkus, B., Bethge, M., Wichmann, F. A., and Brendel, W. (2020). On the surprising similarities between supervised and self-supervised models. arXiv preprint arXiv:2010.08377.

34. Geirhos, R., Narayanappa, K., Mitzkus, B., Thieringer, T., Bethge, M., Wichmann, F. A., and Brendel, W. (2021). Partial success in closing the gap between human and machine vision. Advances in Neural Information Processing Systems, 34:23885–23899.

35. Golan, T., Guo, W., Schütt, H. H., and Kriegeskorte, N. (2022). Distinguishing representational geometries with controversial stimuli: Bayesian experimental design and its application to face dissimilarity judgments. arXiv preprint arXiv:2211.15053.

36. Golan, T., Raju, P. C., and Kriegeskorte, N. (2020). Controversial stimuli: Pitting neural networks against each other as models of human cognition. Proceedings of the National Academy of Sciences, 117(47):29330– 29337.

37. Gower, J. C. (1966). Some distance properties of latent root and vector methods used in multivariate analysis. Biometrika, 53(3-4):325–338.

38. Goyal, P., Duval, Q., Reizenstein, J., Leavitt, M., Xu, M., Lefaudeux, B., Singh, M., Reis, V., Caron, M., Bojanowski, P., Joulin, A., and Misra, I. (2021). VISSL.

39. Goyal, P., Duval, Q., Seessel, I., Caron, M., Singh, M., Misra, I., Sagun, L., Joulin, A., and Bojanowski, P. (2022). Vision models are more robust and fair when pretrained on uncurated images without supervision. arXiv preprint arXiv:2202.08360.

40. Güçlü, U. and van Gerven, M. A. (2015). Deep neural networks reveal a gradient in the complexity of neural representations across the ventral stream. Journal of Neuroscience, 35(27):10005–10014.

41. Güçlü, U. and van Gerven, M. A. (2015). Deep neural networks reveal a gradient in the complexity of neural representations across the ventral stream. Journal of Neuroscience, 35(27):10005–10014. Publisher: Soc Neuroscience.

42. Han, Y., Poggio, T., and Cheung, B. (2023). System identification of neural systems: If we got it right, would we know? arXiv preprint arXiv:2302.06677.

43. Hasson, U., Levy, I., Behrmann, M., Hendler, T., and Malach, R. (2002). Eccentricity bias as an organizing principle for human high-order object areas. Neuron, 34(3):479–490.

44. Hubel, D. H. and Wiesel, T. N. (1968). Receptive fields and functional architecture of monkey striate cortex. The Journal of physiology, 195(1):215–243. Publisher: Wiley Online Library.

45. Jing, Y., Yang, Y., Feng, Z., Ye, J., Yu, Y., and Song, M. (2019). Neural style transfer: A review. IEEE transactions on visualization and computer graphics, 26(11):3365–3385.

46. Kaniuth, P. and Hebart, M. N. (2022). Feature-reweighted representational similarity analysis: A method for improving the fit between computational models, brains, and behavior. NeuroImage, 257:119294.

47. Kanwisher, N., Khosla, M., and Dobs, K. (2023). Using artificial neural networks to ask ‘why’ questions of minds and brains. Trends in Neurosciences.

48. Kaplan, J., McCandlish, S., Henighan, T., Brown, T. B., Chess, B., Child, R., Gray, S., Radford, A., Wu, J., and Amodei, D. (2020). Scaling laws for neural language models. arXiv preprint arXiv:2001.08361.

49. Kay, K. N. (2018). Principles for models of neural information processing. NeuroImage, 180:101–109.

50. Khaligh-Razavi, S.-M., Henriksson, L., Kay, K., and Kriegeskorte, N. (2017a). Fixed versus mixed rsa: Explaining visual representations by fixed and mixed feature sets from shallow and deep computational models. Journal of Mathematical Psychology, 76:184–197.

51. Khaligh-Razavi, S.-M., Henriksson, L., Kay, K., and Kriegeskorte, N. (2017b). Fixed versus mixed RSA: Explaining visual representations by fixed and mixed feature sets from shallow and deep computational models. Journal of Mathematical Psychology, 76:184–197. Publisher: Elsevier.

52. Khaligh-Razavi, S.-M. and Kriegeskorte, N. (2014). Deep supervised, but not unsupervised, models may explain IT cortical representation. PLoS computational biology, 10(11). Publisher: Public Library of Science.

53. Konkle, T. and Alvarez, G. A. (2022). A self-supervised domain-general learning framework for human ventral stream representation. Nature Communications, 13(1):1–12. Publisher: Nature Publishing Group.

54. Krakauer, J. W., Ghazanfar, A. A., Gomez-Marin, A., MacIver, M. A., and Poeppel, D. (2017). Neuroscience needs behavior: correcting a reductionist bias. Neuron, 93(3):480–490.

55. Kriegeskorte, N. (2015). Deep neural networks: a new framework for modeling biological vision and brain information processing. Annual review of vision science, 1:417–446. Publisher: Annual Reviews.

56. Kriegeskorte, N., Mur, M., and Bandettini, P. A. (2008a). Representational similarity analysis-connecting the branches of systems neuroscience. Frontiers in systems neuroscience, 2:4. Publisher: Frontiers.

57. Kriegeskorte, N., Mur, M., Ruff, D. A., Kiani, R., Bodurka, J., Esteky, H., Tanaka, K., and Bandettini, P. A. (2008b). Matching categorical object representations in inferior temporal cortex of man and monkey. Neuron, 60(6):1126–1141. Publisher: Elsevier.

58. Krizhevsky, A., Sutskever, I., and Hinton, G. E. (2012). ImageNet classification with deep convolutional neural networks. In Advances in Neural Information Processing Systems, pages 1097–1105.

59. Leclerc, G., Ilyas, A., Engstrom, L., Park, S. M., Salman, H., and Mądry, A. (2023). Ffcv: Accelerating training by removing data bottlenecks. In Proceedings of the IEEE/CVF Conference on Computer Vision and Pattern Recognition, pages 12011–12020.

60. LeCun, Y., Bengio, Y., and Hinton, G. (2015). Deep Learning. nature, 521(7553):436–444. Publisher: Nature Publishing Group.

61. Lee, H., Margalit, E., Jozwik, K. M., Cohen, M. A., Kanwisher, N., Yamins, D. L., and DiCarlo, J. J. (2020). Topographic deep artificial neural networks reproduce the hallmarks of the primate inferior temporal cortex face processing network. bioRxiv, pages 2020–07.

62. Lin, T.-Y., Maire, M., Belongie, S., Hays, J., Perona, P., Ramanan, D., Dollár, P., and Zitnick, C. L. (2014). Microsoft coco: Common objects in context. In European conference on computer vision, pages 740–755. Springer.

63. Linsley, D., Rodriguez, I. F., Fel, T., Arcaro, M., Sharma, S., Livingstone, M., and Serre, T. (2023). Performance-optimized deep neural networks are evolving into worse models of inferotemporal visual cortex. arXiv preprint arXiv:2306.03779.

64. Liu, Z., Mao, H., Wu, C.-Y., Feichtenhofer, C., Darrell, T., and Xie, S. (2022). A convnet for the 2020s. In Proceedings of the IEEE/CVF Conference on Computer Vision and Pattern Recognition, pages 11976–11986.

65. Long, B., Yu, C.-P., and Konkle, T. (2018). Mid-level visual features underlie the high-level categorical organization of the ventral stream. Proceedings of the National Academy of Sciences, 115(38):E9015–E9024. Publisher: National Acad Sciences.

66. Lüdecke, D., Ben-Shachar, M. S., Patil, I., and Makowski, D. (2020). Extracting, computing and exploring the parameters of statistical models using R. Journal of Open Source Software, 5(53):2445.

67. Mahon, B. Z. and Caramazza, A. (2011). What drives the organization of object knowledge in the brain? Trends in cognitive sciences, 15(3):97–103.

68. Margalit, E., Lee, H., Finzi, D., DiCarlo, J. J., Grill-Spector, K., and Yamins, D. L. (2023). A unifying principle for the functional organization of visual cortex. bioRxiv, pages 2023–05.

69. Marques, T., Schrimpf, M., and DiCarlo, J. J. (2021). Multi-scale hierarchical neural network models that bridge from single neurons in the primate primary visual cortex to object recognition behavior. bioRxiv. Publisher: Cold Spring Harbor Laboratory.

70. McGreivy, N. and Hakim, A. (2022). Convolutional layers are not translation equivariant. arXiv preprint arXiv:2206.04979.

71. Morey, R. D. et al. (2008). Confidence intervals from normalized data: A correction to cousineau (2005). Tutorials in Quantitative Methods for Psychology, 4(2):61–64.

72. Mu, N., Kirillov, A., Wagner, D., and Xie, S. (2021). SLIP: Self-supervision meets Language-Image Pre-training. arXiv preprint arXiv:2112.12750.

73. Muggeo, V. M. (2003). Estimating regression models with unknown break-points. Statistics in medicine, 22(19):3055–3071.

74. Muggeo, V. M. (2008). segmented: an r package to fit regression models with broken-line relationships. R News, 8(1):20–25.

75. Muggeo, V. M. (2017). Interval estimation for the breakpoint in segmented regression: a smoothed score-based approach. Australian New Zealand Journal of Statistics, 59:311–322.

76. Muttenthaler, L., Dippel, J., Linhardt, L., Vandermeulen, R. A., and Kornblith, S. (2022). Human alignment of neural network representations. arXiv preprint arXiv:2211.01201.

77. Naseer, M. M., Ranasinghe, K., Khan, S. H., Hayat, M., Shahbaz Khan, F., and Yang, M.-H. (2021). Intriguing properties of vision transformers. Advances in Neural Information Processing Systems, 34:23296–23308.

78. Nayebi, A., Kong, N. C., Zhuang, C., Gardner, J. L., Norcia, A. M., and Yamins, D. L. (2021). Unsupervised Models of Mouse Visual Cortex. bioRxiv. Publisher: Cold Spring Harbor Laboratory.

79. Nonaka, S., Majima, K., Aoki, S. C., and Kamitani, Y. (2021). Brain hierarchy score: Which deep neural networks are hierarchically brain-like? IScience, 24(9):103013.

80. Olshausen, B. A., Field, D. J., and others (1995). Sparse coding of natural images produces localized, oriented, bandpass receptive fields. Submitted to Nature. Available electronically as ftp://redwood.psych.cornell.edu/pub/papers/sparse-coding.ps. Publisher: Citeseer.

81. Op de Beeck, H. P., Haushofer, J., and Kanwisher, N. G. (2008). Interpreting fmri data: maps, modules and dimensions. Nature Reviews Neuroscience, 9(2):123–135.

82. Paszke, A., Gross, S., Massa, F., Lerer, A., Bradbury, J., Chanan, G., Killeen, T., Lin, Z., Gimelshein, N., Antiga, L., Desmaison, A., Kopf, A., Yang, E., DeVito, Z., Raison, M., Tejani, A., Chilamkurthy, S., Steiner, B., Fang, L., Bai, J., and Chintala, S. (2019). PyTorch: An Imperative Style, High-Performance Deep Learning Library. In Wallach, H., Larochelle, H., Beygelzimer, A., Alché-Buc, F. d., Fox, E., and Garnett, R., editors, Advances in Neural Information Processing Systems 32, pages 8024–8035. Curran Associates, Inc.

83. Pedersen, T. L. (2022). ggforce: Accelerating ’ggplot2’. https://ggforce.data-imaginist.com, https://github.com/thomasp85/ggforce.

84. Pedregosa, F., Varoquaux, G., Gramfort, A., Michel, V., Thirion, B., Grisel, O., Blondel, M., Prettenhofer, P., Weiss, R., Dubourg, V., Vanderplas, J., Passos, A., Cournapeau, D., Brucher, M., Perrot, M., and Duchesnay, E. (2011). Scikit-learn: Machine Learning in Python. Journal of Machine Learning Research, 12:2825–2830.

85. Popham, S. F., Huth, A. G., Bilenko, N. Y., Deniz, F., Gao, J. S., Nunez-Elizalde, A. O., and Gallant, J. L. (2021). Visual and linguistic semantic representations are aligned at the border of human visual cortex. Nature Neuroscience, 24(11):1628–1636. Publisher: Nature Publishing Group.

86. Prince, J. S., Charest, I., Kurzawski, J. W., Pyles, J. A., Tarr, M. J., and Kay, K. N. (2022). Improving the accuracy of single-trial fMRI response estimates using GLMsingle. Elife, 11:e77599. Publisher: eLife Sciences Publications Limited.

87. Prince, J. S. and Konkle, T. (2020). Computational evidence for integrated rather than specialized feature tuning in category-selective regions. Journal of Vision, 20(11):1577–1577.

88. Prince, J. S. and Konkle, T. (2023). Lesioning category-selective units in silico yields functionally specialized deficits. Vision Sciences Society.

89. Puigcerver, J., Riquelme, C., Mustafa, B., Renggli, C., Pinto, A. S., Gelly, S., Keysers, D., and Houlsby, N. (2020). Scalable transfer learning with expert models. arXiv preprint arXiv:2009.13239.

90. R Core Team (2013). R: A Language and Environment for Statistical Computing. R Foundation for Statistical Computing, Vienna, Austria. ISBN 3-900051-07-0.

91. Radford, A., Kim, J. W., Hallacy, C., Ramesh, A., Goh, G., Agarwal, S., Sastry, G., Askell, A., Mishkin, P., Clark, J., and others (2021). Learning transferable visual models from natural language supervision. In International conference on machine learning, pages 8748–8763. tex.organization: PMLR arXiv preprint arXiv:2103.00020.

92. Raghu, M., Unterthiner, T., Kornblith, S., Zhang, C., and Dosovitskiy, A. (2021). Do vision transformers see like convolutional neural networks? Advances in Neural Information Processing Systems, 34:12116–12128.

93. Ratan Murty, N. A., Bashivan, P., Abate, A., DiCarlo, J. J., and Kanwisher, N. (2021). Computational models of category-selective brain regions enable high-throughput tests of selectivity. Nature communications, 12(1):5540.

94. Ren, Y. and Bashivan, P. (2023). How well do models of visual cortex generalize to out of distribution samples? bioRxiv, pages 2023–05.

95. Richards, B. A., Lillicrap, T. P., Beaudoin, P., Bengio, Y., Bogacz, R., Christensen, A., Clopath, C., Costa, R. P., de Berker, A., Ganguli, S., and others (2019). A deep learning framework for neuroscience. Nature Neuroscience, 22(11):1761–1770. Publisher: Nature Publishing Group US New York.

96. Ridnik, T., Ben-Baruch, E., Noy, A., and Zelnik-Manor, L. (2021). Imagenet-21k pretraining for the masses. arXiv preprint arXiv:2104.10972.

97. Rust, N. C. and Movshon, J. A. (2005). In praise of artifice. Nature neuroscience, 8(12):1647–1650.

98. Sax, A., Emi, B., Zamir, A. R., Guibas, L. J., Savarese, S., and Malik, J. (2018). Mid-Level Visual Representations Improve Generalization and Sample Efficiency for Learning Visuomotor Policies.

99. Sax, A., Zhang, J. O., Emi, B., Zamir, A., Savarese, S., Guibas, L., and Malik, J. (2019). Learning to Navigate Using Mid-Level Visual Priors. *arXiv:1912.11121 [cs]*. arXiv: 1912.11121.

100. Schrimpf, M., Kubilius, J., Hong, H., Majaj, N. J., Rajalingham, R., Issa, E. B., Kar, K., Bashivan, P., Prescott-Roy, J., Geiger, F., Schmidt, K., Yamins, D. L. K., and DiCarlo, J. J. (2018a). Brain-Score: Which Artificial Neural Network for Object Recognition is most Brain-Like? bioRxiv preprint.

101. Schrimpf, M., Kubilius, J., Hong, H., Majaj, N. J., Rajalingham, R., Issa, E. B., Kar, K., Bashivan, P., Prescott-Roy, J., Geiger, F., Schmidt, K., Yamins, D. L. K., and DiCarlo, J. J. (2018b). Brain-Score: Which Artificial Neural Network for Object Recognition is most Brain-Like? bioRxiv preprint.

102. Schrimpf, M., Kubilius, J., Lee, M. J., Murty, N. A. R., Ajemian, R., and DiCarlo, J. J. (2020). Integrative benchmarking to advance neurally mechanistic models of human intelligence. Neuron, 108(3):413–423. Publisher: Elsevier.

103. Serre, T. (2019). Deep learning: the good, the bad, and the ugly. Annual Review of Vision Science, 5:399–426. Publisher: Annual Reviews.

104. Sorscher, B., Ganguli, S., and Sompolinsky, H. (2022a). Neural representational geometry underlies few-shot concept learning. Proceedings of the National Academy of Sciences, 119(43):e2200800119.

105. Sorscher, B., Geirhos, R., Shekhar, S., Ganguli, S., and Morcos, A. (2022b). Beyond neural scaling laws: beating power law scaling via data pruning. Advances in Neural Information Processing Systems, 35:19523– 19536.

106. St-Yves, G. and Naselaris, T. (2018). The feature-weighted receptive field: an interpretable encoding model for complex feature spaces. NeuroImage, 180:188–202.

107. Storrs, K. R., Kietzmann, T. C., Walther, A., Mehrer, J., and Kriegeskorte, N. (2021). Diverse Deep Neural Networks All Predict Human Inferior Temporal Cortex Well, After Training and Fitting. Journal of Cognitive Neuroscience, 33(10):2044–2064. Publisher: MIT Press One Rogers Street, Cambridge, MA 02142-1209, USA journals-info . . . .

108. Tang, J., Du, M., Vo, V. A., Lal, V., and Huth, A. G. (2023). Brain encoding models based on multimodal transformers can transfer across language and vision. arXiv preprint arXiv:2305.12248.

109. Tarhan, L. and Konkle, T. (2020). Reliability-based voxel selection. NeuroImage, 207:116350. Publisher: Elsevier.

110. Team, T. M. M. (2021). composer. https://github.com/mosaicml/composer/.

111. Wang, A., Tarr, M., and Wehbe, L. (2019). Neural taskonomy: Inferring the similarity of task-derived representations from brain activity. Advances in Neural Information Processing Systems, 32.

112. Wang, A. Y., Kay, K., Naselaris, T., Tarr, M. J., and Wehbe, L. (2022). Incorporating natural language into vision models improves prediction and understanding of higher visual cortex. BioRxiv, pages 2022–09. Publisher: Cold Spring Harbor Laboratory.

113. Wang, Z., Bovik, A. C., Sheikh, H. R., and Simoncelli, E. P. (2004). Image quality assessment: from error visibility to structural similarity. IEEE transactions on image processing, 13(4):600–612.

114. Wen, H., Shi, J., Chen, W., and Liu, Z. (2018). Deep residual network predicts cortical representation and organization of visual features for rapid categorization. Scientific reports, 8(1):1–17. Publisher: Nature Publishing Group.

115. Wightman, R. (2019). Pytorch image models. https://github.com/rwightman/pytorch-image-models.

116. Wightman, R., Touvron, H., and Jégou, H. (2021). Resnet strikes back: An improved training procedure in timm. arxiv 2021. arXiv preprint arXiv:2110.00476.

117. Willeke, K. F., Fahey, P. G., Bashiri, M., Pede, L., Burg, M. F., Blessing, C., Cadena, S. A., Ding, Z., Lurz, K.-K., Ponder, K., and others (2022). The Sensorium competition on predicting large-scale mouse primary visual cortex activity. arXiv preprint arXiv:2206.08666.

118. Wood, J. N., Lee, D., Wood, B., and Wood, S. M. (2020). Reverse engineering the origins of visual intelligence. In CogSci.

119. Wortsman, M., Ilharco, G., Kim, J. W., Li, M., Kornblith, S., Roelofs, R., Lopes, R. G., Hajishirzi, H., Farhadi, A., Namkoong, H., et al. (2022). Robust fine-tuning of zero-shot models. In Proceedings of the IEEE/CVF Conference on Computer Vision and Pattern Recognition, pages 7959–7971.

120. Wu, Y., Kirillov, A., Massa, F., Lo, W.-Y., and Girshick, R. (2019). Detectron2.

121. Xiao, W. and Kreiman, G. (2020). XDream: Finding preferred stimuli for visual neurons using generative networks and gradient-free optimization. PLoS computational biology, 16(6):e1007973. Publisher: Public Library of Science San Francisco, CA USA.

122. Yamins, D. L. and DiCarlo, J. J. (2016). Using goal-driven deep learning models to understand sensory cortex. Nature neuroscience, 19(3):356. Publisher: Nature Publishing Group.

123. Yamins, D. L., Hong, H., Cadieu, C. F., Solomon, E. A., Seibert, D., and DiCarlo, J. J. (2014). Performance-optimized hierarchical models predict neural responses in higher visual cortex. Proceedings of the National Academy of Sciences, 111(23):8619–8624. Publisher: National Acad Sciences.

124. Yerxa, T., Kuang, Y., Simoncelli, E., and Chung, S. (2023). Learning efficient coding of natural images with maximum manifold capacity representations. arXiv preprint arXiv:2303.03307.

125. Yun, S., Han, D., Oh, S. J., Chun, S., Choe, J., and Yoo, Y. (2019). Cutmix: Regularization strategy to train strong classifiers with localizable features. In Proceedings of the IEEE/CVF international conference on computer vision, pages 6023–6032.

126. Zamir, A. R., Sax, A., Shen, W., Guibas, L. J., Malik, J., and Savarese, S. (2018). Taskonomy: Disentangling task transfer learning. In Proceedings of the IEEE Conference on Computer Vision and Pattern Recognition, pages 3712–3722.

127. Zhang, H., Cisse, M., Dauphin, Y. N., and Lopez-Paz, D. (2017). mixup: Beyond empirical risk minimization. arXiv preprint arXiv:1710.09412.

128. Zhou, H.-Y., Lu, C., Yang, S., and Yu, Y. (2021). ConvNets vs. Transformers: Whose visual representations are more transferable? In Proceedings of the IEEE/CVF International Conference on Computer Vision, pages 2230–2238.

129. Zhuang, C., Yan, S., Nayebi, A., Schrimpf, M., Frank, M. C., DiCarlo, J. J., and Yamins, D. L. (2021). Unsupervised neural network models of the ventral visual stream. Proceedings of the National Academy of Sciences, 118(3). Publisher: National Acad Sciences.

